# An autonomous AI agent for universal behavior analysis

**DOI:** 10.1101/2025.05.15.653585

**Authors:** Almir Aljović, Zuwan Lin, Wenbo Wang, Xinhe Zhang, Arnau Marin-Llobet, Ningyue Liang, Bradley Canales, Jaeyong Lee, Jongmin Baek, Ren Liu, Catherine Li, Na Li, Jia Liu

## Abstract

Behavior analysis across species represents a fundamental challenge in neuroscience, psychology, and ethology, typically requiring extensive expert knowledge and labor-intensive processes that limit research scalability and accessibility. We introduce BehaveAgent, an autonomous multimodal AI agent designed to automate behavior analysis from video input without retraining or manual intervention. Unlike conventional methods that require manual behavior annotation, video segmentation, task-specific model training, BehaveAgent leverages the reasoning capabilities of multimodal large language models (LLM) to generalize across novel behavioral domains without need for additional training. It integrates LLMs, vision-language models (VLMs), and large-scale visual grounding modules, orchestrated through a multimodal context memory and goal-directed attention mechanism, to enable robust zero-shot visual reasoning across species and experimental paradigms, including plants, insects, rodents, primates, and humans. Upon receiving a video input, BehaveAgent autonomously identifies the correct analysis strategy and performs end-to-end behavior analysis and interpretation without human supervision. Leveraging vision-language representations, it performs general-purpose tracking, pose estimation and segmentation. We demonstrate BehaveAgent’s universal applicability to autonomously (1) identify the behavioral paradigm and develop an action plan specialized for the identified paradigm, (2) identify relevant subjects and objects, (3) track those features, (4) identify behavioral sequences with explicit reasoning, (5) generate and execute code for targeted analysis and (6) generate comprehensive research reports that integrate behavioral findings with relevant scientific literature. Through interpretable agentic reasoning, BehaveAgent makes its internal decision-making process transparent, clarifying why particular features are tracked or behaviors inferred. By reducing the time and expertise required for behavior analysis, BehaveAgent introduces a scalable, generalizable, and explainable paradigm for advancing biological and behavioral research.

## Main

Behavior across species represents one of the most complex outputs of the nervous system, serving as the primary window into cognitive, emotional, and social processes. Understanding behavior is fundamental to a wide range of scientific fields, including neuroscience, psychology, pharmacology, and animal welfare, yet behavior analysis presents significant challenges^1-3^. Traditional approaches require labor intensive manual definition of behaviors, behavior coding, training or fine-tuning supervised models for feature tracking, and data interpretation by human experts^4-6^. These approaches are not only time consuming but also limited by subjectivity and inconsistency across different analysts or studies^7^, limiting their accessibility, scalability and generalizability.

Recent technological advancements have greatly increased our capacity to capture rich behavioral data^8^ through high-speed cameras and pose-estimation tools^9, 10^. Despite their precise and detailed feature tracking, these systems depend on users manually labeling a subset of frames to fine-tune models^11^. Furthermore, once fine-tuned for a specific context, such as a particular species or experimental setup, these models often fail to generalize to new environments and experimental contexts, requiring additional retraining and manual labeling. These models are fundamentally limited by their ability to only learn fixed input-output mappings for predefined tasks, limiting their potential use for building generalizable behavior AI agents capable of analyzing diverse behavior experiments and paradigms.

Current computational methods for behavior analysis also lack the adaptive, agentic capabilities needed to autonomously perceive, reason, and take independent actions when analyzing complex behavioral data. Unlike human analysts who adaptively interpret behaviors based on context, recognize novel patterns, and adjust their analytical approach in real time, current computational approaches operate within rigid, predefined frameworks. This limitation prevents them from engaging in flexible hypothesis generation, autonomous exploration of behavioral patterns, or integration of contextual factors that influence animal behavior.

The emergence of multimodal large language models (LLMs)^12^ and vision-language models (VLMs)^13^ has created new possibilities for AI systems that can understand and reason about complex visual data. These models, trained on vast datasets of text and images, demonstrate emergent capabilities^14^ in visual understanding^15^, contextual reasoning^16^, and natural language generation^17^. Once embedded in autonomous AI agents^18, 19^, they are capable of continuously perceiving their environment, making decisions, and acting toward defined goals, offering a promising approach to behavior analysis. AI agents built on multimodal LLMs have exhibited powerful emergent capabilities, including advanced reasoning abilities, contextual interpretation, and strong potential for zero-shot or few-shot generalization^20^, enabling them to adapt to novel data and analytical tasks with minimal supervision. These capacities have already enabled breakthroughs in fields such as neural computation^21^, spatial biology^22^, and chemistry^23^, highlighting their potentials for facilitating scientific discovery^24^.

Here, we introduce BehaveAgent, the first fully autonomous multimodal AI agent for universal behavior analysis. The BehaveAgent integrates multimodal LLMs, VLMs, and visual grounding^25, 26^ techniques, orchestrated by goal-oriented agentic reasoning and supported by a multimodal context memory system. This integration enables a continuous analytical workflow, where BehaveAgent can autonomously (1) identify the behavioral paradigm and develop specialized objectives for the analysis, (2) identify relevant subjects and objects, (3) track relevant features, (4) identify behavioral sequences and provide explicit reasoning, and (5) generate comprehensive research reports that integrate behavioral findings with relevant scientific literature.

BehaveAgent demonstrates explainable reasoning capabilities, which provides detailed insights into its decision-making. This enables transparency and interpretability of the entire analytical process, which allows researchers to validate conclusions within their experimental frameworks. BehaveAgent also demonstrates broad generalization across diverse biological contexts from plants to humans without requiring specialized training. By autonomously identifying and annotating behavioral events, it eliminates time intensive manual behavioral registration, the need for training or fine-tuning new models, thereby allowing researchers to focus on higher level data interpretation and hypothesis testing.

Importantly, BehaveAgent consolidates the generated multimodal data into comprehensive, structured deep research reports that transform raw behavioral data into actionable scientific insights. These reports not only document the analytical process and findings but also integrate relevant scientific literature, interpret the observed behaviors in their biological context, and suggest directions for future experiments.

In addition to operating in autonomous mode, BehaveAgent also supports human-AI collaboration, assisting researchers in addressing specific questions, operationally defining behaviors, and suggesting, designing and executing appropriate analyses.

By providing a unified approach for analyzing behavior across species and contexts, BehaveAgent enables researchers to investigate behavioral questions at greater scale and consistency, with potential to advance how behavior is studied and understood.

## Results

### BehaveAgent architecture and adaptive core

The conventional workflow of behavior analysis (**Fig. 1a**) requires extensive manual labor, including time-consuming expert labeling, feature extraction, and model training for each specific behavioral context. This process is not only resource intensive but also suffers from poor generalizability across different experimental paradigms. Traditional methods typically target behavior and coding experts as their intended users, requiring manual data labeling and training resources, resulting in low generalizability.

**Fig. 1:**
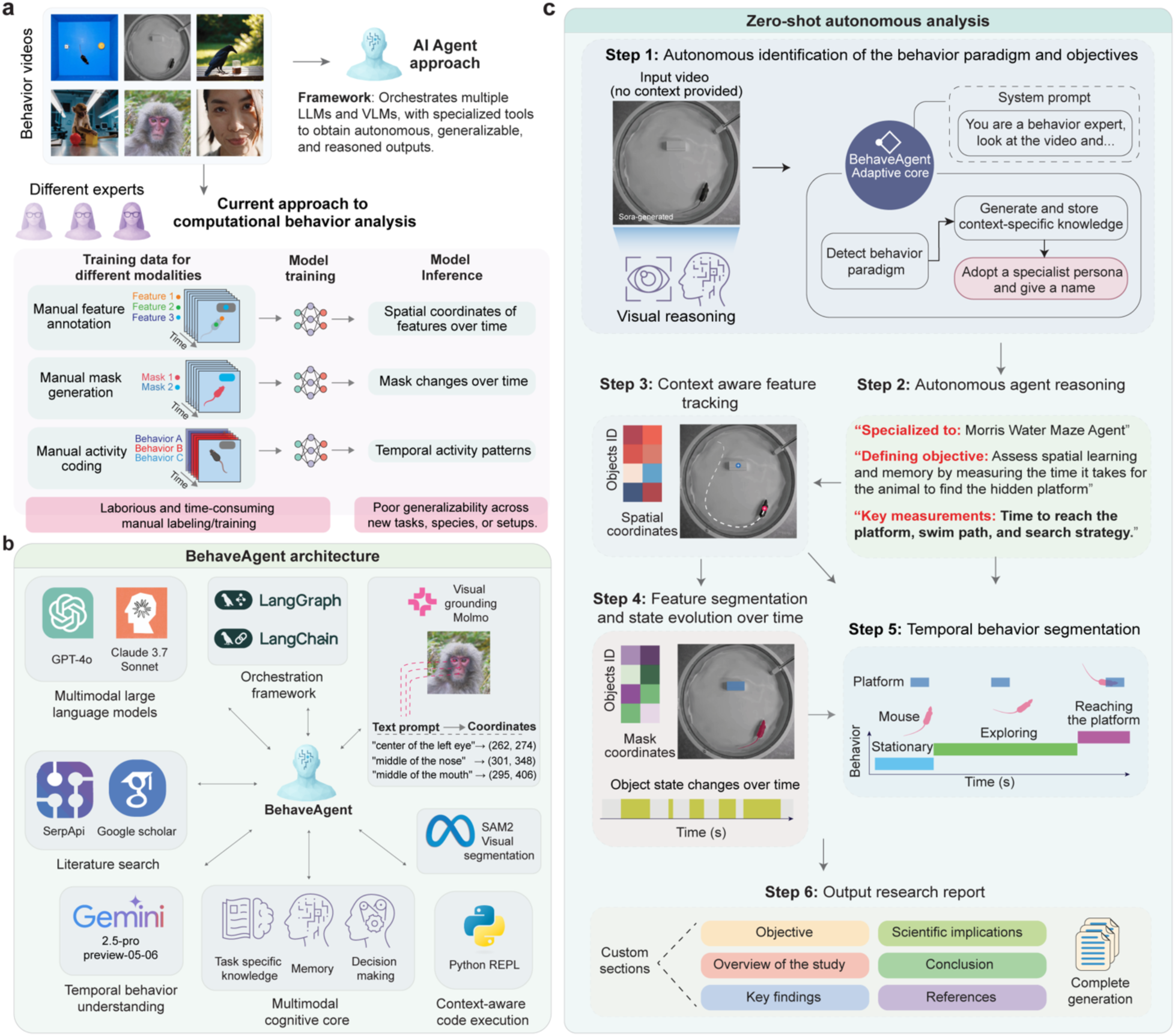
BehaveAgent architecture. (**a**) Comparison between current methodology for behavior analysis and the AI Agent method. Traditional approaches require manual expert labeling, time-consuming feature extraction, and resource intensive model training, resulting in poor generalizability across experimental contexts. In contrast, BehaveAgent offers an automatic, time efficient framework with interpretable outputs accessible to users without specialized expertise. (**b**) BehaveAgent’s integrated architecture combines AI agent orchestration frameworks (LangGraph, LangChain), visual processing models (visual grounding, segmentation), temporal segmentation, dynamic code execution capabilities, literature search functionality (SerpApi, Google Scholar), multimodal large language models (GPT-4o, Claude 3.7 Sonnet), and a multi-modal cognitive core that integrates task-specific knowledge, memory, and decision-making processes. (**c**) Illustration of the agent’s ability to function autonomously and generate analysis without prior training or context. BehaveAgent’s adaptive core activates upon receiving an input video, performing video reasoning to identify the behavioral context and automatically transforming into a specialized agent. The specialized agent then proceeds through a systematic workflow: defining the experimental type and objective (Step 1), engaging in autonomous reasoning to determine key measurements (Step 2), performing feature tracking with spatial coordinates (Step 3), conducting feature segmentation (Step 4), generating temporal behavior segmentation (Step 5), and producing a comprehensive research report (Step 6). Sora-generated refers to the use of AI for video generation.

In contrast, BehaveAgent’s architecture (**Fig. 1b**) is built around a central autonomous AI agent that orchestrates multiple AI components through an integrated framework. At its core, BehaveAgent leverages multimodal LLMs such as GPT-4o^27^, Claude-3.7-Sonnet^28^ and Gemini-2.5-Pro-Preview-05-06^29^ for their reasoning and planning capabilities essential for behavioral analysis. These models are complemented by specialized visual processing components, including large visual grounding models such as Molmo^26^ that enable precise object localization and tracking using natural language instructions that can be generated by LLMs, Segment Anything Model 2 (SAM2)^30^ for visual segmentation, and Gemini-2.5-Pro-Preview-05-06 for temporal behavior understanding. BehaveAgent integrates with literature search capabilities through services such as Google Scholar and SerpApi^31^, allowing the agent to autonomously incorporate relevant scientific literature into its analysis and ground behavioral interpretations in current peer-reviewed research. The AI agent orchestration framework, such as LangGraph and LangChain^32^, coordinates these components and manages information flow, enabling a multi-modal cognitive core that integrates task-specific knowledge, memory systems, and decision-making processes. Additionally, BehaveAgent can dynamically generate and execute Python code^33^ tailored to specific analytical requirements, eliminating the need for manual programming.

BehaveAgent’s fully autonomous mode (**Fig. 1c; Supplementary Video 1**) initiates with zero-shot behavior paradigm detection and autonomous goal setting, enabled by its adaptive component, we refer to as adaptive core. This process identifies the relevant behavior paradigms within the video input and formulates specialized action plans for analysis. Specifically, even without prior contextual information, BehaveAgent can autonomously interpret the content of a video to determine the most appropriate analytical approach. For instance, it can recognize that a given video captures the *Morris Water Maze* paradigm for rodent behavior and proceed with a self-generated, context-specific analysis plan, entirely without human intervention or additional training. This capability is driven by a sophisticated visual reasoning system that seamlessly integrates vision-language models (VLMs) with contextual understanding, enabling it to interpret diverse behavioral scenarios across various species. These large-scale pretrained VLMs encode general visual concepts and relationships, empowering BehaveAgent to achieve universal video scene comprehension, including the recognition of experimental setups, subjects, and behavioral patterns, without requiring domain-specific fine-tuning.

After identifying the behavior paradigm, the BehaveAgent can make a systematic analytical workflow: (i) identifying the behavioral paradigm and its objectives, (ii) determining the appropriate analysis approach, and (iii) performing context-aware feature tracking, feature segmentation, and temporal behavior segmentation. Ultimately, this process generates a comprehensive, context-aware analysis report that integrates visual analysis with relevant scientific literature, providing a complete research document with objectives, findings, implications, methodology, and references.

Our results showed that zero-shot behavior paradigm detection and autonomous goal setting is the key to enable the accurate autonomous downstream behavior analysis. Therefore, to test this capability of BehaveAgent, we utilized AI-generated behavioral videos created by OpenAI’s Sora text-to-video model, which provide a controlled yet diverse set of behavioral scenarios. When presented with these Sora-generated videos without any contextual information (**Fig. 2a**), BehaveAgent automatically analyzes the content, employing frame sequence sampling and VLM understanding to identify the behavioral paradigm present in each video (**Fig. 2b**). For example, BehaveAgent successfully identifies four distinct behavioral paradigms and automatically sets analysis objectives, renaming itself based on the identified paradigm: a *Morris Water Maze Agent* for rodent spatial learning assessment (Video 1), a *Corvid Problem-Solving Agent* for avian cognition analysis (Video 2), a *Primate Object Manipulation Agent* for studying non-human primate cognitive abilities (Video 3), and a *Facial Expression Analysis Agent* for human emotion research (Video 4).

**Fig. 2:**
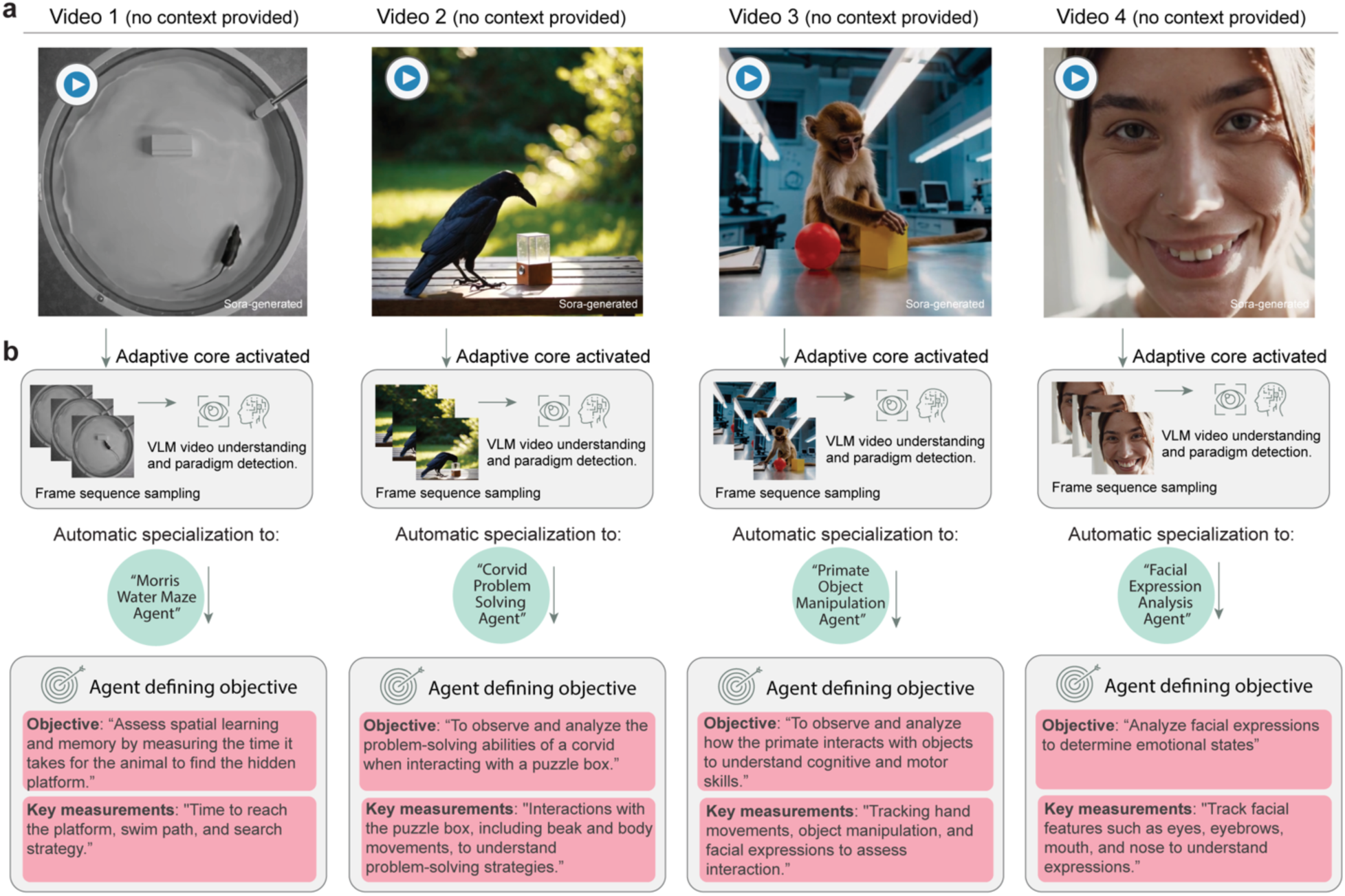
Zero-shot behavior paradigm detection and goal setting using BehaveAgent. (**a**) Diverse videos generated by OpenAI’s Sora text-to-video model presented to BehaveAgent without any contextual information: a rodent in a Morris Water Maze (Video 1), a corvid interacting with a puzzle box (Video 2), a primate manipulating objects (Video 3), and a human displaying facial expressions (Video 4). (**b**) For each video, BehaveAgent’s adaptive core activates automatically, performing frame sequence sampling and vision-language model (VLM) understanding to identify the behavioral paradigm present in the video. Without any prior training on these specific contexts, BehaveAgent autonomously transforms into the appropriate specialized agent for each scenario: a Morris Water Maze Agent for rodent spatial learning assessment, a Corvid Problem-Solving Agent for avian cognition analysis, a Primate Object Manipulation Agent for studying non-human primate cognitive abilities, and a Facial Expression Analysis Agent for human emotion research. For each specialized configuration, BehaveAgent autonomously defines appropriate research objectives and key measurements tailored to the specific behavioral context. Sora-generated refers to the use of AI for video generation.

For each video analysis, BehaveAgent also autonomously define appropriate research objectives and key measurements tailored to the specific context (**Fig. 2b**). For the Morris Water Maze video, the system identifies that the objective is to “assess spatial learning and memory by measuring the time it takes for the animal to find the hidden platform” with key measurements including “time to reach the platform, swim path, and search strategy.” For the corvid problem-solving task, the objective becomes “to observe and analyze the problem-solving abilities of a corvid when interacting with a puzzle box” with measurements focusing on “interactions with the puzzle box, including beak and body movements.” Similarly, the primate object manipulation analysis focuses on the objective as “to observe and analyze how the primate interacts with objects to understand cognitive and motor skills” and identifies key measurements including “tracking hand movements, object manipulation, and facial expressions to assess interaction.” while human facial analysis focuses on “analyze facial expressions to determine emotional states” with key measurements including “track facial features such as eyes, eyebrows, mouth, and nose to understand expressions.”

These results show that we can efficiently test and validate BehaveAgent’s performance across diverse scenarios, species, and contexts, showing its versatility and reliability. Without prior training on specific behavioral paradigms, BehaveAgent autonomously and accurately identifies diverse experiments and establishes appropriate analytical frameworks, eliminating the need for manual configuration and reducing technical barriers to advanced behavioral analysis across multiple research domains.

After recognizing behavior paradigm and making the action plan, BehaveAgent can move forward with the context-aware feature tracking. Here, a key technological innovation enabling BehaveAgent’s universal applicability to autonomously perform the context-aware feature tracking is the implementation of visual grounding models such as Molmo^25^ (**Extended Data Fig. 1**). Conventional methods require manual labeling of features, task-specific model training, and fine-tuning for each new context, resulting in non-generalizable models across different species or experimental paradigms (**Extended Data Fig. 1**a). In contrast, BehaveAgent uses visual grounding models, which are pretrained on large-scale image-text pairs, capable of understanding natural language descriptions and their visual correspondences without requiring domain-specific fine-tuning (**Extended Data Fig. 1**b). Guided by natural language prompts generated from the previous behavior paradigm identification and action plan development step, it allows for autonomous context-aware feature tracking in BehaveAgent.

Specifically, this visual grounding system processes input images through a vision encoder and connector that interface with an LLM. As an example, when the LLM generates a prompt such as “*Point to the subject’s nose*”, the visual grounding models can precisely locate features across diverse contexts without requiring new training data or model adjustments, allowing efficient transitions between tracking different features across species and contexts and autonomous behavior video analysis.

Furthermore, our results showed that this integrated architecture could enable BehaveAgent to function “out of the box” across diverse species and behavioral paradigms, from rodent maze navigation to plant behavior and human facial expression analysis. The system’s ability to autonomously adapt to new contexts without additional training represents a major advancement in behavior analysis technology, offering researchers a powerful tool that combines the flexibility of human analysis with the efficiency and consistency of automated systems.

### Goal-oriented feature detection and reasoning-based tracking

To demonstrate its autonomous performance, we recorded a mouse performing a novel object recognition test, a standard test in neuroscience studying cognitive functions in mice. The video, given an unidentifiable name, was provided to the agent without any additional context.

After inputting this video (**Fig. 3a**), BehaveAgent performed frame sampling and VLM-based video understanding to identify and store context-dependent information. This initial processing allows the BehaveAgent to understand the experimental setup without requiring any instructions from the user. The system autonomously identifies the behavior paradigm, in this case, a Novel Object Recognition Test, and defines the objective: “*To measure a rodent’s exploratory behavior and preference between familiar and novel objects, assessing recognition memory and cognitive function.*” Key measurements are identified as “*time spent investigating each object, frequency of approaches to each object, movement patterns of the rodent throughout the arena*”.

**Fig. 3:**
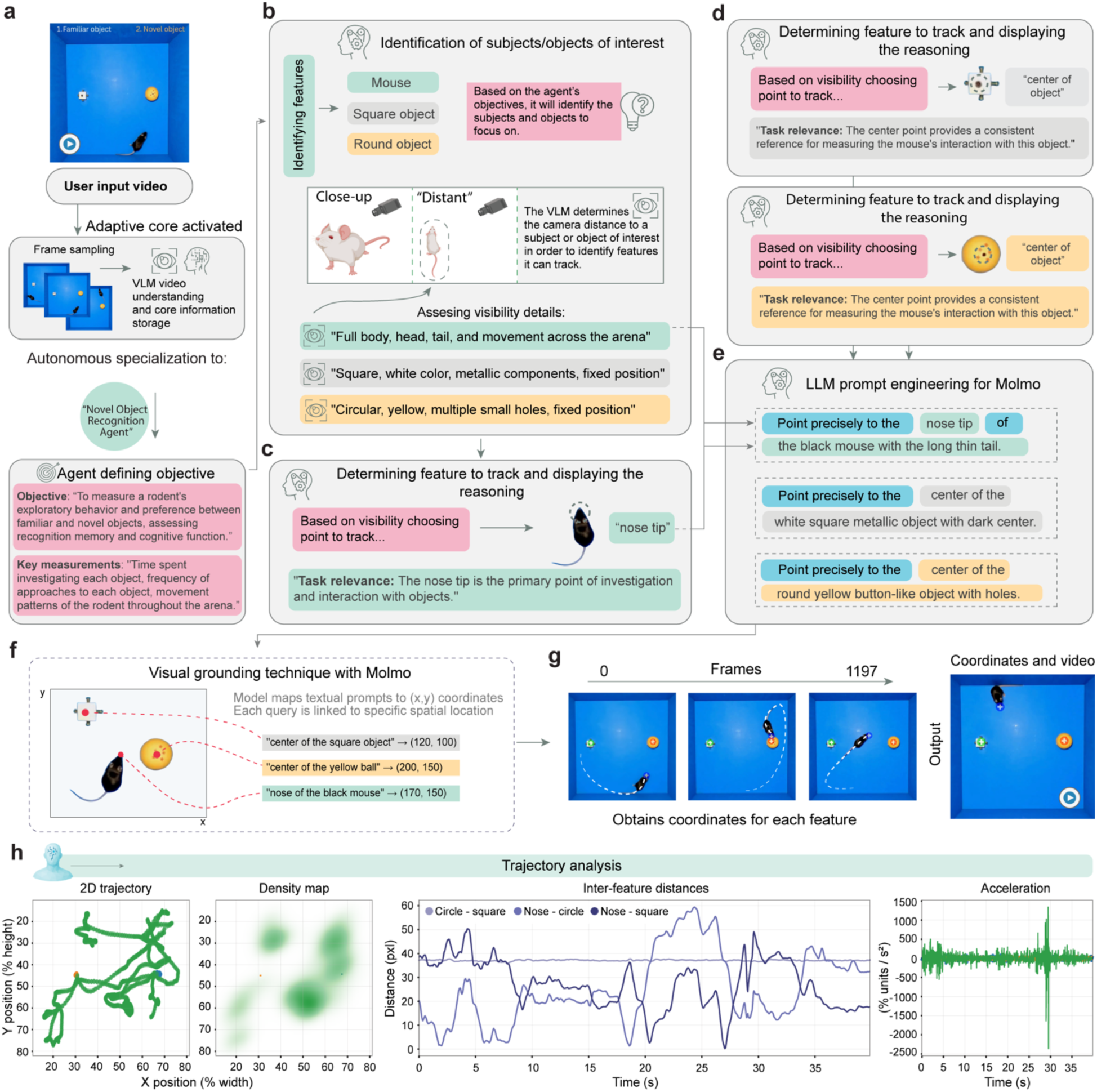
Goal-oriented feature detection and reasoning-based tracking. (**a**) Upon receiving a user input video showing a novel object recognition experiment, BehaveAgent’s adaptive core activates, performing frame sampling and VLM-based video understanding. The system automatically transforms into a Novel Object Recognition Agent and defines the objective as assessing the animal’s ability to recognize a novel object compared to a familiar one. (**b**) BehaveAgent identifies subjects and objects of interest based on the agent’s objectives and assesses visibility details for each element. The system also determines the camera distance for each subject/object to inform tracking decisions. (**c-d**) For each identified element, BehaveAgent determines the optimal tracking point based on visibility and task relevance, providing explicit reasoning for each decision. For example, it chooses to track the mouse’s nose because “tracking the nose helps determine which object the rat is interacting with and for how long” and selects the center points of both objects to measure proximity and interaction duration. (**e**) BehaveAgent uses LLM prompt engineering for Molmo, generating specific textual prompts that pinpoint exact features to track. (**f**) Schematics showing that natural language prompts are processed through Molmo, which map the textual descriptions to exact (x,y) coordinates in the video frames. (**g**) The resulting coordinate tracking enables BehaveAgent to follow the position of each feature across all video frames, creating a comprehensive spatial record of the mouse’s movements and interactions with both objects. (**h**) This coordinate data is then used for SAM2, a video segmentation model, which segments and tracks the objects throughout the video sequence.

BehaveAgent then proceeds through a systematic reasoning process to identify subjects and objects of interest as well as camera settings (**Fig. 3b; Supplementary Video 2**). Based on the agent’s objectives, it identifies three key features: the mouse, a square object, and a round object. The system assesses visibility details for each element, noting specific characteristics such as “*full body, head, tail, and movement across the arena*” for the mouse, “*square, white color, metallic components, fixed position*” for the square object, and “*circular, yellow, multiple small holes, fixed position*” for the round object. The system also automatically determines camera distance to subjects, distinguishing between close-up and distant views to optimize its tracking approach.

Next, BehaveAgent explicitly reasons about which features to track (**Fig. 3c-d**), which is the critical aspect of the autonomous tracking approach. For each identified element, the system determines the optimal tracking point based on both visibility considerations and task relevance. This dual-criteria approach ensures that the tracking is both technically feasible and scientifically meaningful. For example, it chooses to track the mouse’s “*nose tip*” because it is “*the primary point of investigation and interaction with objects,*” a decision directly tied to the experimental objective of assessing object recognition. Similarly, it selects the center points of both objects to measure proximity and interaction duration.

To implement precise tracking, BehaveAgent autonomously uses LLM prompt engineering for Molmo visual grounding capacities (**Fig. 3e**), generating specific textual prompts that pinpoint features to track. These prompts are natural language descriptions such as “*Point precisely to the nose tip of the black mouse with the long thin tail*,” These natural language prompts are processed through visual grounding techniques with Molmo (**Fig. 3f**), which maps the textual descriptions to exact (x,y) coordinates in the video frames. This visual grounding capability represents a crucial bridge between language-based reasoning and precise spatial localization.

The resulting coordinate tracking (**Fig. 3g**) enables BehaveAgent to follow the position of each feature across all video frames, creating a comprehensive spatial record of the mouse’s movements and interactions with both objects. This continuous tracking provides the foundation for quantitative analysis of the animal’s behavior, including measurements of time spent near each object and patterns of exploration. BehaveAgent then performed trajectory analysis (**Fig. 3h**), generating multiple analytical visualizations including 2D trajectories showing the mouse’s movement patterns, density maps revealing spatial preferences, inter-feature distance plots measuring proximity between the mouse and objects over time, and acceleration data capturing the dynamics of movement. These visualizations transform raw coordinate data into scientifically meaningful representations that directly address the experimental objectives.

To evaluate BehaveAgent’s tracking accuracy, we applied it to an Open Field test (**Extended Data Fig. 2**a-c) compared to human annotators (**Extended Data Fig. 2**d, e). 3D trajectory plots created from data by human annotators and BehaveAgent show a high level of similarity, confirming that, for this specific task, BehaveAgent achieves tracking accuracy comparable to that of human experts.

In brief, we demonstrate that BehaveAgent’s key characteristic is its use of explicit reasoning at each step of the process. The results show that rather than simply applying pretrained tracking models, BehaveAgent dynamically and autonomously determines what to track based on experimental objectives and provides clear explanations for these decisions. This goal-oriented approach ensures that tracking is directly relevant to the behavioral analysis task, while comprehensive reasoning makes the entire process transparent and interpretable.

### Object segmentation and VLM-guided temporal state tracking

Building on the precise coordinate tracking established previously, we further implemented sophisticated object segmentation and feature clustering to analyze how subjects and objects change over time. This capability is essential for understanding complex behavioral interactions, where object states may change due to subject manipulation.

BehaveAgent begins the segmentation process by passing the previously identified context-aware object coordinates to the SAM2 model (**Fig. 4a**). Using the (x,y) coordinates obtained through visual grounding from the previous step, BehaveAgent then uses SAM2 to segment and track each identified object throughout the video sequence. This process results in precise masks for each object across all frames. This enables detailed analysis of shape changes and interactions, creating a foundation for analyzing how elements interact and change over time.

**Fig. 4:**
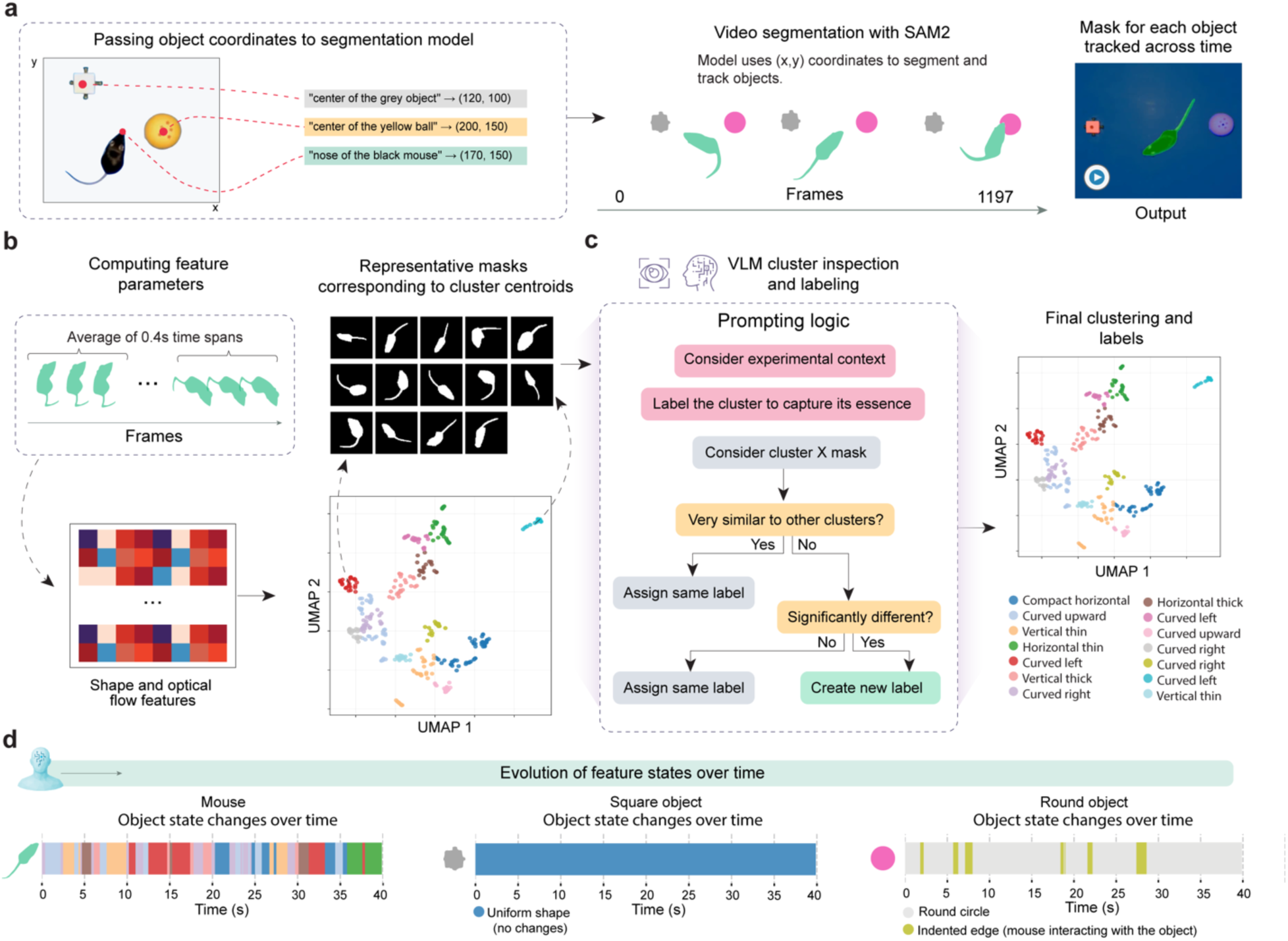
Video object segmentation and feature clustering with VLM-guided semantic labeling for temporal state tracking. **(a)** Schematic representation of the interaction between Molmo’s visual grounding and SAM2 video object segmentation. The obtained (x, y) coordinates are fed into the SAM2 segmentation model, enabling consistent object masks and tracking across the video. **(b)** Schematic of feature extraction and embedding results. First, for each segmented object, shape descriptors and optical-flow features were computed over short (0.4 s) time spans and averaged to obtain high-dimensional feature vectors. These vectors were then projected into two dimensions via UMAP, revealing visually separable clusters whose centroids were represented by exemplar masks. **(c)** Logic of VLM-guided cluster inspection and semantic labeling. **(d)** Temporal state tracking. VLM-derived labels were mapped back onto each frame to annotate object states over time. Shown are representative state-sequence plots for each object in the experiment.

To capture temporal dynamics of object states, we further implemented a feature clustering approach in BehaveAgent (**Fig. 4b**). The system computes feature parameters by analyzing short time spans throughout the video, extracting shape and optical flow features that capture both static appearance and dynamic movement characteristics. These features are dimensionally reduced using UMAP, creating a 2D representation, where similar object states cluster together. For each cluster, BehaveAgent generates representative masks corresponding to the cluster centroids, providing visual examples of each state.

Here, another key innovation in BehaveAgent’s approach is its use of VLM-guided semantic labeling to interpret and annotate these clusters (**Fig. 4c**). Rather than treating clusters as arbitrary groupings, BehaveAgent uses robust prompting logic that considers the experimental context when assigning meaningful labels. The system examines each cluster’s representative masks, determines their similarity to other clusters, and makes informed decisions about whether to assign existing labels or create new ones. This context-aware labeling process transforms abstract visual patterns into scientifically meaningful categories that directly relate to the behavioral analysis objectives.

Specifically, to ensure comprehensive coverage of potential object states while maintaining interpretability, BehaveAgent initially implements overclustering (**Extended Data Fig. 3**). This approach deliberately creates more clusters than might be strictly necessary, capturing fine-grained differences in object states. The VLM then inspects these clusters, identifying which represent genuinely distinct states versus minor variations of the same state. For example, **Extended Data Fig. 3**a shows how multiple clusters of the round object are consolidated into two main states: “*Round Center*” (normal state) and “*Indented Edge*” (when the mouse interacts with it). Similarly, **Extended Data Fig. 3**b demonstrates how the square object clusters are all recognized as representing a single “*Uniform Shape*” state, indicating no significant changes over time.

The final output of these integrated processes is a temporal mapping of object states throughout the video (**Fig. 4d**). BehaveAgent generates timelines showing how the mouse and each object transition between different states over the course of the experiment. For the mouse, this reveals a complex sequence of behavioral states as it explores and interacts with objects. The square object maintains a uniform shape throughout, while the round object alternates between its normal state and an “*indented edge*” state during mouse interactions.

Implementing this segmentation and state tracking capability enables BehaveAgent to capture subtle behavioral interactions that would be difficult to identify through coordinate tracking alone. By combining precise segmentation with VLM-guided semantic interpretation, the system transforms raw visual data into meaningful behavioral insights, revealing patterns that might otherwise remain hidden in complex video data.

### Context-aware temporal behavior segmentation

Temporal behavior segmentation represents one of the most challenging aspects of behavioral analysis, traditionally requiring extensive human expertise to manually identify meaningful behavioral transitions and patterns in continuous data streams. We further address this challenge by implementing two context-aware agentic AI approaches that integrate multiple analytical perspectives with explicit reasoning at each step.

Building on the previously established tracking and object state analysis, BehaveAgent aggregates a comprehensive set of multimodal data (**Fig. 5a**) that serves as the foundation for behavior segmentation. This includes experimental context, agent’s goals, raw coordinates, anomaly calculations, speed and acceleration metrics, inter-feature distances, trajectory plots, mask area changes, and mask cluster sequences. BehaveAgent implements a three-stage temporal segmentation logic (**Fig. 5b**) that progressively refines behavioral boundaries and classifications. Stage 1 (Initial assessment) combines contextual information with generated data to perform rule-based segmentation, which is then refined using LLM reasoning. Stage 2 (VLM-based refinement) improves the segmentation through VLM validation by using subset of frames as visual evidence to validate behavior and temporal boundaries. Stage 3 employs semantic understanding to produce context-aware behavior categories.

**Fig. 5:**
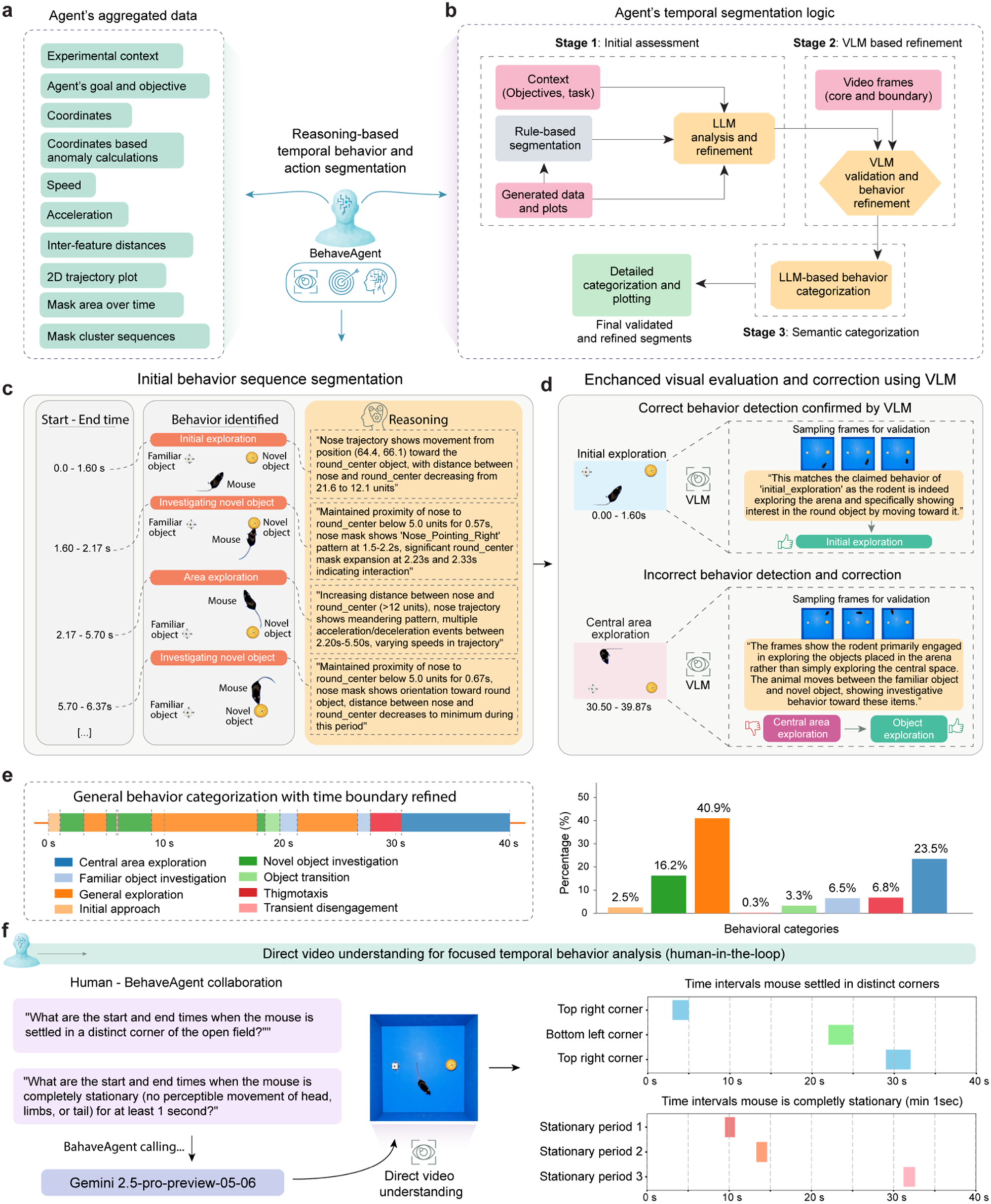
Context-aware behavior temporal behavior segmentation. **(a)** BehaveAgent examining multiple data streams simultaneously, including raw experimental coordinates, calculated metrics (speed, acceleration, inter-feature distances), 2D trajectory plots and mask area and mask patterns over time. **(b)** Schematic representation of temporal segmentation logic. BehaveAgent uses a combination of LLM reasoning and VLM inspection to analyze these data streams, identify behavioral patterns, predict temporal segmentation boundaries, and provide explicit reasoning for its decisions. **(c)** Based on analysis in (**b**), BehaveAgent generating an initial behavior sequence segmentation, dividing the continuous behavioral stream into discrete, meaningful segments with detailed reasoning for each segmentation decision; Transparent insert images are adapted for visualization. **(d)** BehaveAgent-enhanced visual evaluation and correction using VLMs, reviewing its initial behavior detections, confirming correct identifications and correcting any errors. **(e)** Final behavior segmentation and context-aware categorization producing a detailed timeline of behaviors. The general behavior categorization further aggregates the detailed behaviors into broader categories with the associated bar chart quantifying the percentage of time spent in each general behavioral category. (**f**) Human-in-the-loop mode for direct video understanding for focused temporal behavior analysis.

Specifically, the initial behavior sequence segmentation (**Fig. 5c**) divides the continuous behavioral stream into discrete, meaningful segments. Each segment is defined by a start and end time, an identified behavior and a visual representation showing the positions of the rodent and objects. Crucially, BehaveAgent provides detailed reasoning for each segmentation decision, explaining exactly why a particular behavior was identified. For example, it notes that during the “*initial exploration*” segment (0.0-1.60s), “*Nose trajectory shows movement from position (94.4, 66.1) toward the round_center object, with distance between nose and round_center decreasing from 21.9 to 12.1 units.*”. This reasoning is guided by the agent’s initially defined objectives and the experimental context.

To ensure accuracy, BehaveAgent implements enhanced visual evaluation and correction using VLMs (**Fig. 5d**). The system reviews its initial behavior detections by sampling frames from each segment and using VLM analysis to confirm correct identifications or correct errors. When an incorrect behavior detection occurs, such as misclassifying “*object exploration*” as “*central area exploration,*” the VLM provides a correction based on visual evidence: “*The frames show the rodent primarily engaged in exploring the objects placed in the arena rather than simply exploring the central space. The animal moves between the familiar object and novel object, showing investigative behavior toward these items.*”

These validated and corrected behavior detections form the fine-grained classifications shown in the detailed behavior sequence (**Extended Data Fig. 4**a), capturing subtle behaviors such as “*Investigation round object,*” and “*Thigmotaxis exploration.*” To make this detailed analysis more interpretable, BehaveAgent employs semantics-based behavior grouping (**Extended Data Fig. 4**b), where the multimodal LLM uses context-specific knowledge to group these detailed behavior descriptions into broader categories. For example, “*Area exploration*” and “*Extended area exploration*” are merged into the more general category of “*General exploration*.” This hierarchical approach results in a general behavior sequence (**Extended Data Fig. 4**c) that maintains scientific relevance while reducing complexity.

The culmination of this process is the final context-aware temporal segmentation (**Fig. 5e**). BehaveAgent produces a comprehensive timeline of behaviors with refined boundaries, identifying specific actions such as “*Central area exploration*,” “*Novel object investigation*,” “*Thigmotaxis,*” and “*Transient disengagement.*” This detailed segmentation is accompanied by a quantitative analysis showing the percentage of time spent in each behavioral category.

Furthermore, BehaveAgent also offers direct video understanding for focused temporal behavior analysis (**Fig. 5f**). This capability enables human-in-the-loop collaboration, where researchers can ask specific questions about temporal behavior patterns through natural language queries. Examples include “*What are the start and end times when the mouse is settled in a distinct corner of the open field?*” or “*What are the start and end times when the mouse is completely stationary for at least 1 second?*” For this direct video analysis, BehaveAgent uses advanced video understanding capabilities of multimodal LLMs such as Gemini-2.5-Pro-Preview-05-06. The system processes these natural language queries through direct video understanding, generating precise temporal analyses without requiring pre-processed tracking data. The results are presented as timeline visualizations showing, for example, the time intervals when the mouse settled in distinct corners or remained completely stationary. This provides researchers with a flexible, query-based approach to extract specific behavioral information directly from video data.

BehaveAgent’s context-aware temporal segmentation capabilities represent a significant advancement in automated behavior analysis. Unlike conventional methods that rely on arbitrary thresholds or predefined templates, BehaveAgent’s reasoning-based approach considers the temporal dynamics of movements, spatial relationships between subjects and objects, and the specific experimental context. This integrated methodology enables the system to capture subtle transitions between behavioral states and provide explicit reasoning for each identification, either autonomously through its comprehensive analytical pipeline or in response to specific researcher queries through its direct video understanding feature.

### Autonomous research report generation with literature verification

The final stage of BehaveAgent’s analytical pipeline is the generation of comprehensive research reports that integrate behavioral analysis with relevant scholarly literature, transforming raw data into actionable scientific insights. BehaveAgent begins with the multimodal results generated through its analytical pipeline (**Fig. 6a**), including trajectories, inter-feature distances, acceleration profiles, object state changes, and behavioral segmentation timelines. These diverse representations capture different aspects of the behavioral analysis, from spatial patterns to temporal dynamics.

**Fig. 6:**
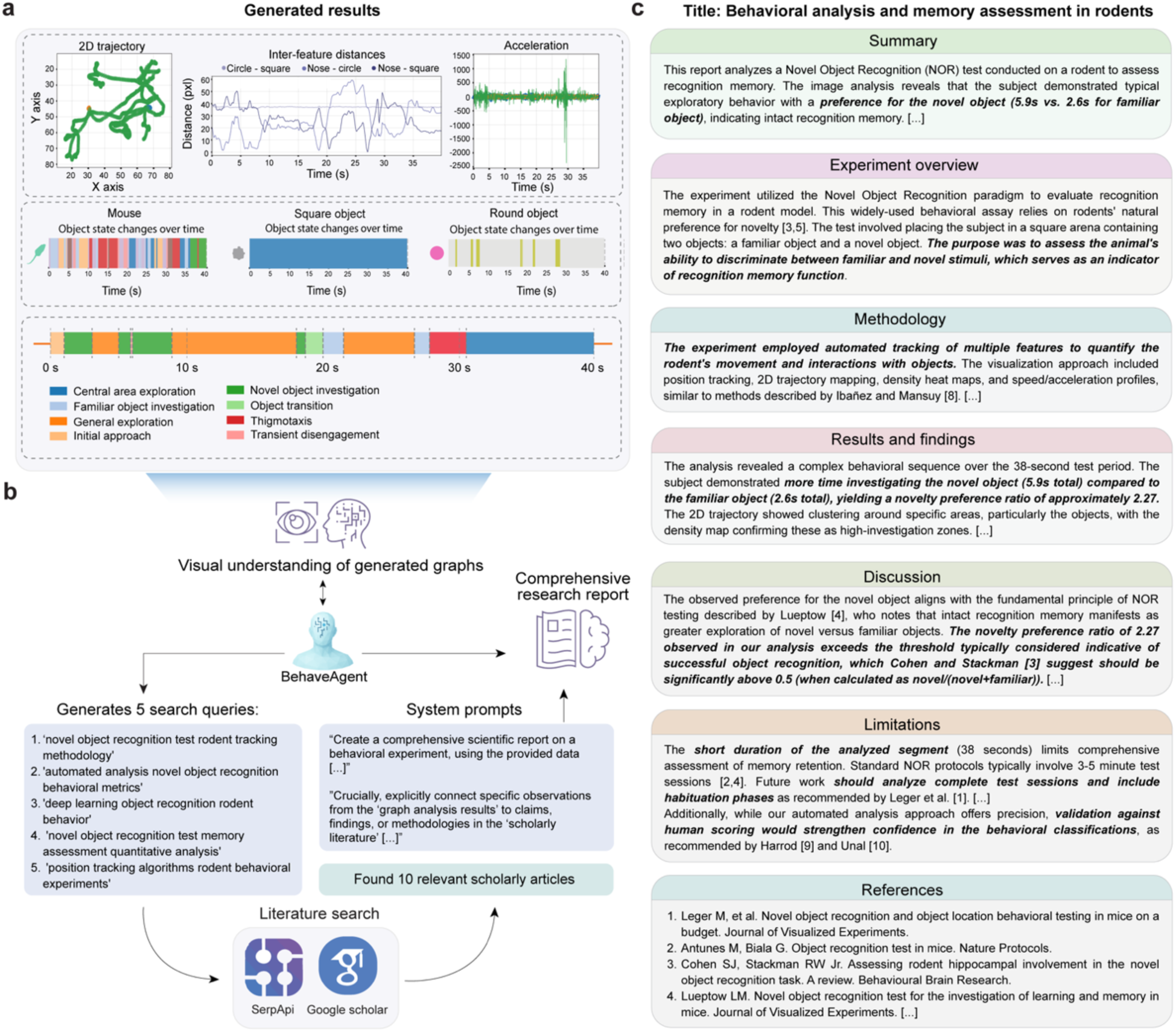
Comprehensive report generation and self-limitation assessment. (**a**) Table showing key metrics that BehaveAgent generated and compiled in previous steps. **(b)** BehaveAgent integrating visual understanding of graphical results with scientific literature to generate a comprehensive behavioral report. The system formulates context-aware search queries, performs automated literature searches (via SerpApi and Google Scholar) to verify the results, and uses LLM prompting to extract and connect relevant findings. **(c)** Generated research report summarizing key behavioral insights from the NOR experiment. The report includes a structured breakdown of the experimental summary, methodology, results, discussion, and limitations. Quantitative observations, such as the rodent’s increased preference for the novel object and calculated preference ratio, are contextualized with references to prior studies from the retrieved literature. The limitations section includes a self-assessment, where the agent recommends comparing its behavior classifications against human annotations to further validate its accuracy and robustness.

To transform these technical outputs into meaningful scientific insights, we employ visual understanding and literature integration (**Fig. 6b**). The BehaveAgent analyzes the generated visualizations and autonomously generates relevant search queries such as “*novel object recognition test rodent tracking methodology*” and “*automated analysis novel object recognition behavioral metrics.*” These queries are processed through SerpApi and Google Scholar, retrieving scholarly articles that provide context for interpreting the behavioral data.

The BehaveAgent then integrates the analytical results with insights from scientific literature to generate a comprehensive research report (**Fig. 6c**). This report follows a standard scientific structure, including summary, experiment overview, methodology, results, discussion, limitations, and references sections.

For example, the report for the representative video analysis generally includes the following sections. The summary section provides a concise overview of the key finding: “*The image analysis reveals that the subject demonstrated typical exploratory behavior with a preference for the novel object (5.9s vs. 2.6s for familiar object), indicating intact recognition memory.*” The experiment overview section contextualizes the analysis within the broader scientific framework, explaining that the *Novel Object Recognition* paradigm evaluates recognition memory in rodent models. The methodology section, connecting the analytical approach to established methods, highlights similarities to techniques described in the literature. The results section presents quantitative insights, highlighting that “*The subject demonstrated more time investigating the novel object (5.9s total) compared to the familiar object (2.6s total), yielding a novelty preference ratio of approximately 2.27*”. The discussion section interprets these findings in relation to established scientific knowledge, noting that “*The novelty preference ratio of 2.27 observed in our analysis exceeds the threshold typically considered indicative of successful object recognition, which Cohen and Stackman [3] suggest should be significantly above 0.5*”. Notably, the report also acknowledges limitations, such as the short duration of the analyzed segment compared to standard protocols. Finally, the report concludes with a properly formatted reference section citing relevant literature, providing a scholarly foundation for the analysis and interpretation.

We envision this autonomous report generation capability represents a significant advancement in behavioral analysis, which can transform complex multimodal data into coherent scientific narratives that integrate analytical results with relevant scholarly knowledge. By connecting observed behavioral patterns to established scientific frameworks, we show that BehaveAgent can provide researchers with actionable insights that can directly inform scientific understanding and future experimental design.

### Cross-species generalization across biological contexts

A fundamental challenge in behavioral analysis is developing a system that can generalize across diverse biological contexts without requiring specialized training or configuration for each new domain. Here, we demonstrate that BehaveAgent can address this challenge through its adaptive architecture and universal visual understanding capabilities.

To test BehaveAgent’s generalization capability, we applied the system to three significantly different biological contexts: plant growth responses (**Fig. 7a**), drosophila courtship behavior (**Fig. 7b; Supplementary Video 3**), and primate object manipulation (**Fig. 7c; Supplementary Video 4**). When presented with these diverse videos without contextual information, BehaveAgent’s adaptive core automatically identified the behavior paradigms and developed the analysis objectives.

**Fig. 7:**
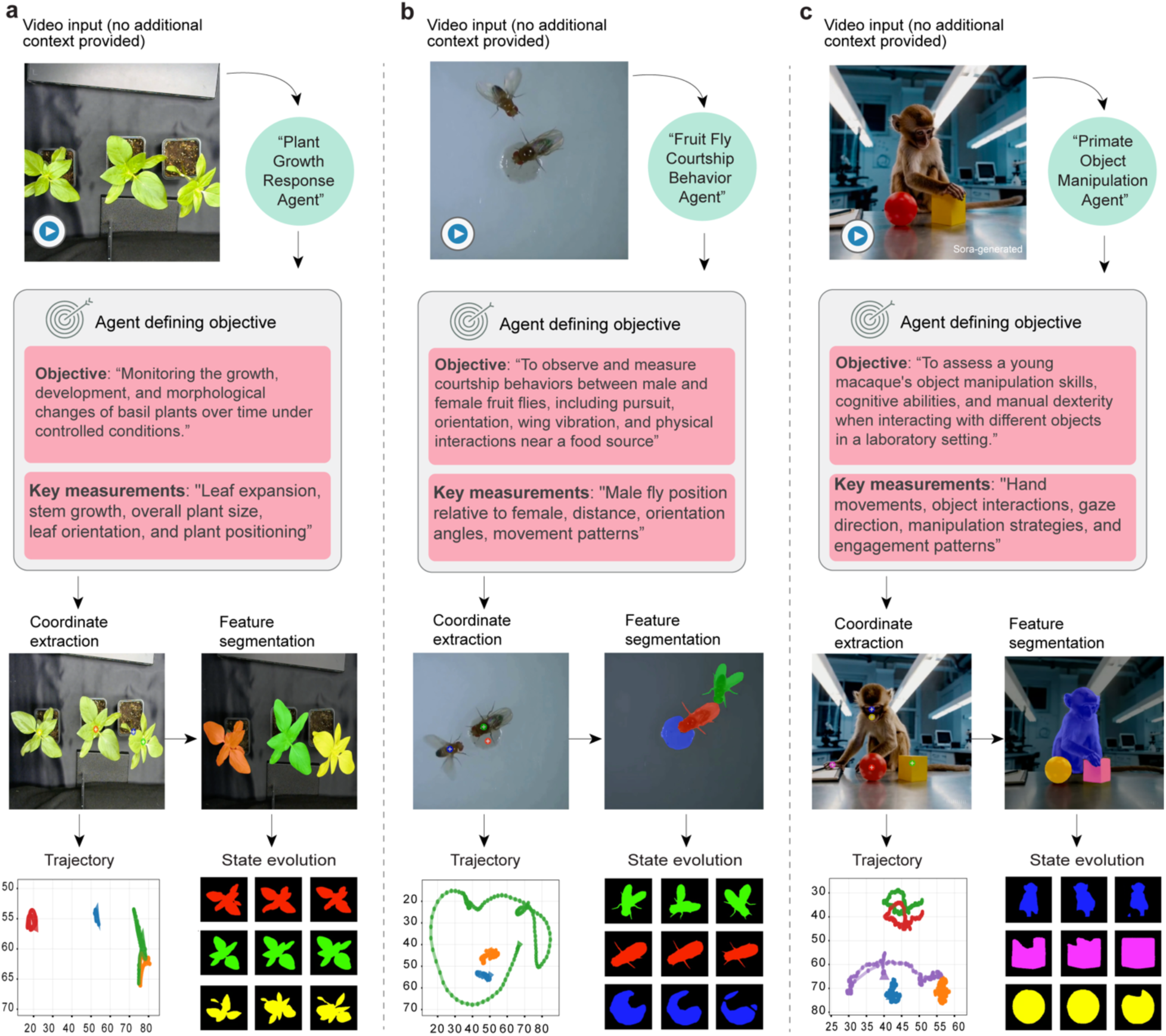
BehaveAgent generalizes across diverse biological contexts. **(a)** BehaveAgent specializing in a “Plant Growth Response Agent”. The agent processes video input of plants to autonomously define its objective: monitoring plant development over time. It extracts coordinates of plant parts, segments features, tracks spatial trajectories, and visualizes state evolution to capture morphological dynamics. The plant video was sourced from ref.^35^. **(b)** BehaveAgent specializing in a “Fruit Fly Courtship Behavior Agent” that analyzes interactions between male and female fruit flies without any prior user-provided context. The agent autonomously identifies courtship behavior as its objective, focusing on male-female positioning and movement patterns. It computes trajectories and generates feature-based segmentations to trace behavioral states over time. The fruit fly video was sourced from ref.^34^. **(c)** BehaveAgent specializing in a “Primate Object Manipulation Agent” that analyzes a juvenile macaque in a laboratory setting and defines its goal as evaluating object manipulation. It identifies hand movements, gaze direction, and manipulation strategies as key features. The agent extracts detailed spatial information, segments body and object features, and maps trajectories and behavioral state transitions. Sora-generated refers to the use of AI for video generation.

For example, for plant behavior analysis (**Fig. 7a**), BehaveAgent autonomously identifies plant behavior as the focus of investigation and renames itself accordingly “*Plant Growth Response Agent*” and defines the objective as “*Monitoring the growth, development, and morphological changes of basil plants over time under controlled conditions*.” The system identifies key measurements including “*Leaf expansion, stem growth, overall plant size, leaf orientation, and plant positioning.*” This demonstrates BehaveAgent’s ability to recognize that plants, despite being non-motile organisms, exhibit analyzable behaviors through growth patterns and morphological changes.

When analyzing fruit fly courtship (**Fig. 7b**), BehaveAgent specializes into a “*Fruit Fly Courtship Behavior Agent*” with the objective “*To observe and measure courtship behaviors between male and female fruit flies, including pursuit, orientation, wing vibration, and physical interactions near a food source.*” The system identifies appropriate measurements for this context, including “*Male fly position relative to female, distance, orientation angles, movement patterns.*”.

For primate behavior analysis (**Fig. 7c**), BehaveAgent detects object manipulation experiment in primates and renames itself to a “*Primate Object Manipulation Agent*” with the objective “*To assess a young macaque’s object manipulation skills, cognitive abilities, and manual dexterity when interacting with different objects in a laboratory setting.*” Key measurements include “*Hand movements, object interactions, gaze direction, manipulation strategies, and engagement patterns,*” reflecting the more complex cognitive and motor behaviors relevant to primate studies.

In each case, BehaveAgent proceeds through the analytical pipeline: coordinate extraction, feature segmentation, trajectory analysis, and state evolution tracking, but adapts the specific implementation details to the unique requirements of each biological context.

Notably, BehaveAgent’s generalization can be easily extended to human facial expression analysis (**Extended Data Fig. 5**). When presented with a Sora-generated video of a person displaying changing facial expressions (**Extended Data Fig. 5**a), BehaveAgent automatically specializes into a “*Facial Expression Analysis Agent.*” The system performs context-aware feature detection (**Extended Data Fig. 5**b), identifying and tracking critical facial landmarks. It conducts context-aware behavior sequence detection (**Extended Data Fig. 5**c), autonomously segmenting the video into distinct emotional states, including “*Neutral expression baseline,*” “*Smile development,*” “*Smile maintenance,*” and “*Sustained smile.*” BehaveAgent also demonstrates robustness to challenging lighting conditions (**Extended Data Fig. 5**d), showing consistent pose-estimation performance across both shadowed and well-lit portions of the face.

We believe that BehaveAgent’s ability to generalize across such diverse biological contexts, from plants to insects to primates to humans, without requiring specialized training for each domain represents a significant advancement in behavior analysis technology. This universal applicability reduces the need for researchers to develop different analytical tools for different experimental paradigms, streamlining the research workflow and enabling more direct comparisons across species and contexts.

### Human-in-the-loop interactive mode for collaborative behavior analysis

In addition to its autonomous capabilities, BehaveAgent has a human-in-the-loop interactive mode, enabling direct collaboration between researchers and the AI. This synergy combines human scientific expertise with AI-driven scientific understanding and analytical power, allowing researchers to address specific questions. The session initiates with BehaveAgent querying the researcher’s goals (**Fig. 8a**), such as experiment type, behaviors of interest, and key research questions. In an example conversation, the researcher details a “*mice social interaction experiment,*” seeking to track both mice, determine interaction initiation and duration, and analyze speed and trajectory changes. In response, BehaveAgent proposes tailored analytical pathways like “*Behavioral Event Analysis*” or “*Detailed Motion Tracking,*” explaining how each aligns with the stated objectives. For instance, using action segmentation for interaction initiation and precise coordinate tracking for trajectory changes for relevant features.

**Figure 8:**
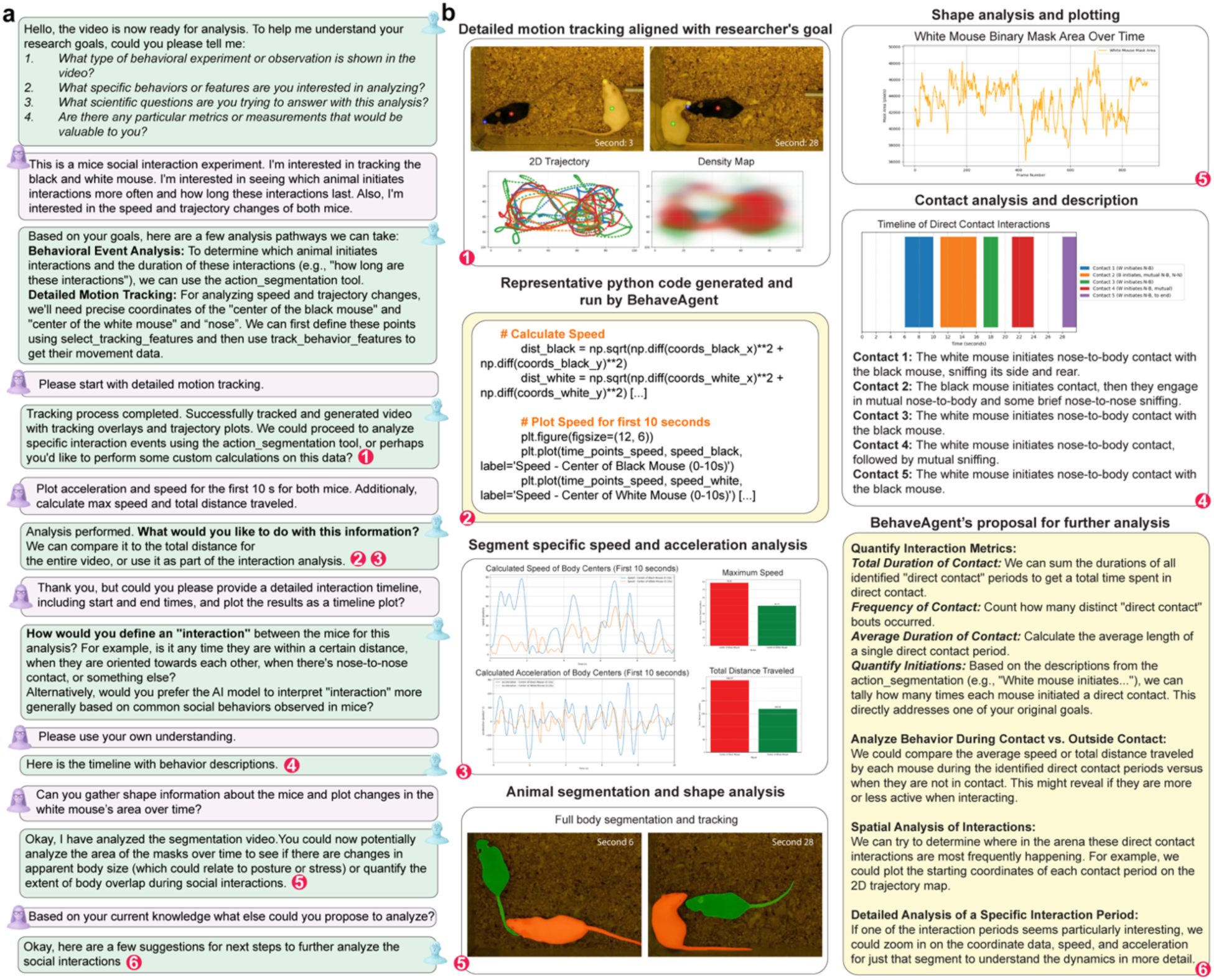
Human-in-the-loop for collaboration with BehaveAgent. **(a)** A sample collaboration between the researcher and BehaveAgent, where the researcher specifies the goals of the analysis, focusing on interaction behaviors between black and white mice. The video was sourced from ref.^37^. BehaveAgent suggests and guides the analysis, utilizing a human-in-the-loop approach, and the researcher provides input and feedback through key prompts, including requests for motion tracking, speed analysis, interaction timelines, and segmentation insights. **(b)** The resulting data outputs, including 2D trajectory maps, speed calculations, body segmentation, and contact timelines, are presented in the interactive chat interface, allowing for dynamic exploration and visualization of the results. This includes code snippets for speed calculation, segment-specific speed plots, and proposals for further analysis.

Upon selecting “*Detailed Motion Tracking*,” BehaveAgent executes the analysis, generating visualizations such as 2D trajectories and density maps (**Fig. 8, panel 1**) that reveal movement patterns and spatial preferences. Researchers can then request specific computations, such as plotting acceleration and speed for an initial segment. BehaveAgent responds by dynamically generating and executing Python code (**Fig. 8, panel 2**), which calculates the required metrics. The resulting segment-specific analyses (**Fig. 8, panel 3**) display speed/acceleration profiles, maximum speed, and total distance. This on-demand code generation democratizes complex analyses and remove manual programming barriers. The system is designed to respect the complexity of behavior analysis, often asking for clarification and operational definitions of behaviors of interest. When queried about defining “*interaction,*” BehaveAgent might suggest a specific operational definition, based on proximity, orientation, contact, or proceed with the model’s understanding of the behavior of interest.

A request for a “*detailed interaction timeline*” prompts BehaveAgent to generate a contact analysis (**Fig. 8, panel 4**), illustrating a precise timeline of distinct events complemented by qualitative descriptions (e.g., “*Contact 1: The white mouse initiates nose-to-body contact…*”). BehaveAgent can also facilitate morphological analyses. If asked to “*gather shape information…and plot changes in the white mouse’s area over time,*” it uses segmentation tool and produce plots of mask area changes (**Fig. 8, panel 5**). Such analysis can reveal subtle postural shifts potentially indicative of emotional or social states. The session can culminate with BehaveAgent suggesting further analytical proposal (**Fig. 8, panel 6**) if prompted. Proposals might include quantifying total contact duration and frequency, comparing behavior during and outside contact, or conducting spatial analysis of interaction hotspots. This proactive guidance aids in developing more sophisticated analytical strategies.

This human-in-the-loop paradigm enables a human-AI collaborative research process. By facilitating scientific dialogue with a range of tools that can be applied in different contexts, BehaveAgent supports the integration of human expertise and AI-driven behavior analysis.

## Discussion

We introduced BehaveAgent, which represents a shift toward a more holistic approach to behavior analysis, offering a fully autonomous and interactive solution that addresses challenges that have long constrained this field. By integrating multimodal LLMs, VLMs, and visual grounding techniques within an orchestrated AI agent framework and memory system, we demonstrated that BehaveAgent can eliminate the need for manual labeling, specialized model training, and context-specific programming, which have traditionally created high barriers to entry in behavioral research. Furthermore, BehaveAgent’s context-driven attention and visual grounding approach offers inherent generalizability through language-based adaptation. This allows the system to precisely locate features across diverse contexts without requiring new training data, simply by adapting its natural language prompts.

BehaveAgent’s explicit reasoning capabilities enhance its scientific value by making the analytical process transparent and interpretable. Unlike traditional machine learning approach, which function as a black box without much interpretability, BehaveAgent offers detailed reasoning for each analytical decision, from why specific features are tracked to how behavioral segments are identified. This transparency builds trust in the system’s outputs and provides researchers with deeper insights into the behavioral patterns being studied. In addition, the system’s hierarchical approach to behavioral categorization balances analytical depth with interpretability, allowing researchers to examine behavior at different levels of granularity. Finally, BehaveAgent’s ability to integrate behavior analysis with relevant scientific literature through autonomous literature searches represents another significant advancement, transforming raw behavioral data into actionable scientific insights contextualized within current research.

While BehaveAgent represents a major advancement, certain limitations should be acknowledged. Most of these limitations stem from the constraints of the underlying models. Currently, we used Molmo as visual grounding technique in BehaveAgent, which offers a single image prompt to coordinate detection. This means that each individual frame of a video is treated independently, as there is no memory of previous states. We attempted to address this issue using various filtering and VLM-based validation methods. Potentially, the agent could rely more on the SAM2 segmentation and tracking output in situations where Molmo demonstrates poor performance, as SAM2 has a memory system that enables consistent object tracking. We also notice that prompt-to-pixel approaches can have limitations in situations where it is difficult for the agent to verbally distinguish between positions that need to be tracked. We see this issue most often in kinematic analysis tasks or when there are multiple objects that appear identical. In its current state, BehaveAgent uses VLM to inspect the features and assigns a unique feature identifier, which is then used to engineer a prompt for Molmo, mitigating some of these issues.

Another limitation is inherent to LLM systems, reflecting their tendency to generate different sequences of words across multiple runs. As a result, users may receive outputs with slightly different wording in multiple runs. While this issue was not observed during visual grounding with Molmo or SAM2, it can also arise in temporal behavior segmentation, affecting both reasoning-based and video LLM approaches. Running the model multiple times and aggregating the outputs, serving as a straightforward form of cross-validation, can help mitigate this variation and yield more robust conclusions.

An additional concern is the computational cost. Since the agent relies on multiple large models, it requires high-performance GPUs to operate efficiently. This challenge can be addressed in the future by integrating BehaveAgent with cloud-based computation and processing systems.

In conclusion, BehaveAgent opens new possibilities for behavioral research by enabling analyses at scales and across contexts previously impractical. The system’s ability to process large volumes of behavioral data without human intervention could reveal patterns that might remain hidden in smaller scale studies. Its analytical approach across different experimental paradigms facilitates more direct comparisons between behavioral studies, potentially leading to more generalizable insights about behavior across species.

We envision that by democratizing access to sophisticated behavior analysis tools and enabling more comprehensive, efficient, and reproducible studies, BehaveAgent has the potential to fundamentally transform how we study behavior across different scientific domains. As AI agent technologies advance, systems like BehaveAgent represent the beginning of a new era where autonomous AI agents serve as collaborative partners in scientific discovery, accelerating our understanding of complex behavioral outputs across diverse organisms.

## Methods

### BehaveAgent framework

We built BehaveAgent on LangChain and LangGraph frameworks, integrating multiple LLMs with advanced vision language models to enhance behavior analysis. The architecture implements a multi-stage analysis pipeline with five sequential processing modules: (1) experiment identification using frame extraction and multimodal LLM understanding, (2) resolution evaluation with feature detection optimization, (3) feature tracking leveraging Molmo’s prompt to pixel capabilities, (4) temporal behavior segmentation, and (5) video object segmentation utilizing Molmo extracted coordinates passed to SAM2 for segmentation. Molmo integration enables direct vision language processing that bypasses the need for training or fine-tuning supervised models for position estimation and tracking. LangGraph provides the stateful workflow architecture necessary for maintaining experimental context across analytical stages, while LangChain facilitates tool integration and memory management. The system employs a cyclic processing workflow with shared memory structures for inter-module data exchange. We implemented human-in-the-loop capabilities with feedback integration through LangChain’s agent framework. The technical implementation utilizes OpenAI API integration, Ultralytics’ computer vision libraries, and AllenAI’s Molmo model (Molmo-7B-D-0924) for visual grounding tasks. BehaveAgent processes standard video formats and outputs quantitative pose vectors, temporally segmented behavioral sequences, and object segmentation masks to facilitate comprehensive behavioral phenotyping in behavior research.

### Adaptive core

In the center of BehaveAgent is its adaptive core that automatically specializes analysis pathways based on experiment specific content. The system first implements experiment type identification by extracting 5 frames at equidistant time points throughout the video for balanced sampling. These frames, along with their relative positions, are analyzed by GPT-4o VLM using a zero-temperature multimodal classification approach with the prompt: “*Analyze these video frames from a behavioral experiment*” followed by specific analytical objectives. The system stores this contextual understanding in a persistent memory architecture with timestamp indexing, enabling retrieval of analysis context in subsequent processing stages. A critical adaptive component dynamically determines specialized agent configurations through the specialized function, which prompts GPT-4o to generate experiment specific agent designations (e.g., “Morris Water Maze Agent”, “Social Interaction Agent”) based on the initial analysis. The prompt specifically instructs: “*determine a specific, descriptive name for an AI agent that would specialize in analyzing this type of experiment*” with formatting requirements to ensure specificity to exact experiment types rather than general categories. This specialized agent designation then drives subsequent feature selection, tracking parameter optimization, and analytical focus. This architecture enables BehaveAgent to automatically adapt its analytical approach to the specific constraints and requirements of each experimental paradigm, providing targeted behavior quantification without requiring manual experiment classification or feature selection.

### Adaptive core benchmarking using AI-generated experimental videos

To evaluate BehaveAgent’s adaptive core capabilities, we implemented a synthetic benchmarking protocol using OpenAI’s Sora generative video model. We engineered distinct text prompts designed to create 5-10 second videos across two categories: laboratory behavioral paradigms (e.g., Morris water maze, primate social interaction), naturalistic environments (e.g., field observations, problem solving in birds, facial expressions in humans), and scenarios with challenging lighting conditions. These AI-generated videos provided ground truth experimental conditions with controlled variability, enabling systematic assessment of the system’s experiment classification accuracy, subject/object detection robustness, and feature selection appropriateness. The synthetic dataset allowed us to evaluate the adaptive core’s performance across diverse experimental paradigms without requiring extensive real world data collection, while maintaining precise control over experimental variables.

### Visual grounding approach

We built BehaveAgent’s visual grounding system on AllenAI’s Molmo models, enabling direct mapping from natural language descriptions to pixel coordinates in behavioral videos. By leveraging Molmo’s multimodal capabilities, we established a prompt to pixel methodology that bypasses the need for supervised training or fine-tuning on labeled datasets. We first employ the GPT-4o or Claude 3.7 Sonnet VLM to analyze sample video frames, automatically determining subjects and objects in the experimental context, which then informs prompt generation for precise feature localization. We engineered a structured prompt template that follows the format “*Point to [precise anatomical description] of the [subject]*” with conditional logic for occlusion handling. This approach generates absolute pixel coordinates across video frames to enable tracking continuity. For deterministic coordinate extraction, we configured Molmo inference with temperature=0.0, max_tokens=100, top_k=1, and seed=42, ensuring completely reproducible outputs across repeated queries. We implemented coordinate extraction through regex-based parsing of XML-formatted responses, with three post-processing approaches for trajectory refinement: median filtering for outlier removal, Kalman filtering for predictive smoothing, and Savitzky-Golay filtering for noise reduction while preserving signal characteristics. Multiple frames were processed concurrently using Python’s asyncio library. A configurable concurrency limit was applied to manage the number of simultaneous requests to the visual language model, optimizing throughput while maintaining the original temporal sequence of the frame data. Default concurrency limit is 50. We deployed this visual grounding infrastructure on a single NVIDIA A100 (40GB SXM4) GPU with 30 vCPUs and 200 GB RAM. The vLLM server hosts either Molmo-7B-D-0924 or Molmo-72B-0924 models, enabling efficient integration with the larger BehaveAgent framework.

### Behavior recordings and videos

Behavior tests were conducted in an open-field environment consisting of a square blue acrylic chamber (40 × 40 cm) with open top. We positioned a GoPro HERO13 Black Creator Edition camera 70 cm above the chamber floor to capture the entire behavioral arena from an overhead perspective. Two objects were affixed to the chamber floor in fixed positions to serve as environmental stimuli. Three-month-old male mice were individually introduced to the testing arena. Mice were allowed to freely explore the environment and interact with the stimulus objects for the entire recording duration. A novel or familiar state of the object in the video was randomly assigned to test the agent’s capacity to make goals and plan analysis based on the experimental context. For additional context testing, we used videos of Drosophila courtship behavior, sunflower growth and mice social interaction from publicly available sources^34, 35, 37^.

### Tool integration

BehaveAgent’s tool integration layer operates around four core AI frameworks: OpenAI GPT-4o, Anthropic Claude-3.7-Sonnet and Google Gemini-2.5-Pro-Preview-05-06 for images, videos, long context analysis, and multimodal reasoning, AllenAI’s Molmo for zero-shot visual grounding, and SAM2 for video object segmentation. Each tool is implemented as a Python class adhering to LangChain’s Tool schema, registered through factory functions that configure callable instances. The ExperimentIdentifier tool implements frame extraction using OpenCV (cv2.VideoCapture) with strategic sampling of 5 frames for temporal coverage. These frames are processed through GPT-4o using its multimodal API with a structured neuroscience expert prompt and deterministic generation parameters (temperature=0.0, max_tokens=512). For extended contextual reasoning and report generation, we integrate Claude-3.7-Sonnet through Anthropic’s API with context window optimization and specialized system prompts for scientific writing. The PoseTracker tool integrates Molmo through a custom API wrapper with parameter settings temperature=0.0, max_tokens=100, top_k=1, and seed=42, enabling reproducible coordinate extraction. For object segmentation, we implemented the SimpleSAM wrapper class utilizing SAM2VideoPredictor with configuration parameters conf=0.25, task=‘segment’, mode=‘predict’, imgsz=1024. The system generates and executes Python code dynamically through a PythonREPLTool and ExecutableTool. This enables runtime generation of custom analyses based on extracted trajectories, with code execution secured through input validation and restricted imports. For literature integration, we implemented a ScholarTool using the Scholarly API (scholarly.search_pubs) with regex pattern matching to extract citation markers from analysis text and insert relevant citations from Google Scholar. These tools are orchestrated through a central agent that maintains execution state in a shared memory context, accessed through structured paths that isolate tool specific data while enabling access to data across tools.

### Temporal behavior segmentation

For temporal behavior segmentation, BehaveAgent uses two approaches: a reasoning-based method and a method that uses the novel Gemini-2.5-pro-preview-05.06 model for video understanding. For the reasoning-based temporal analysis, BehaveAgent operates through a multi-stage pipeline that integrates aggregated data analysis with vision–language-model verification. The behavior segmentation process begins with kinematic pattern extraction from tracking coordinates, identifying behavioral transition points through acceleration analysis, velocity profiling, and spatial positioning. The system employs parallel detection algorithms targeting ethologically relevant behaviors: approach sequences, object investigations, and high-speed movements, generating candidate behavioral segments that undergo contextual validation to remove artifacts and merge related behaviors.

For coherent behavioral narrative construction, we employ Claude-3.7-Sonnet, providing tracking data, mask changes over time, position summaries, and acceleration profiles as contextual input to generate fine-grained classifications with temporal boundaries. We enhanced detection accuracy through vision-language model integration, implementing a frame-based verification system that extracts temporal slices at behavior transitions and submits representative frames to GPT-4o or Claude-3.7-Sonnet for visual confirmation. This process employs specialized prompts that contextualize the putative behavior, with confidence scores weighted based on initial prediction and visual agreement.

Our boundary refinement mechanism implements precise behavior onset/offset detection by extracting equidistant frames within a ±1.0 second window around each candidate transition point. These frames undergo VLM analysis using specialized prompts to identify exact transition frames, calculating temporal correction factors. For ambiguous cases, the system implements a reconciliation workflow that queries both trajectory and visual characteristics with explicit conflict resolution prompts.

The segmentation pipeline generates hierarchical behavior categorization, mapping detailed behaviors to broader scientific categories while maintaining temporal relationships. For visualization, the system creates comprehensive timeline plots showing behavior transitions, durations, and confidence metrics. For the second approach, which is based on VLM video understanding, we utilized the Gemini-2.5-pro-preview-05-06 model. The BehaveAgent agent employs this model through its “action_segmentation” tool, which processes raw video and user or agent-defined prompts to identify the timing of specific actions. For model parameters, we used a temperature of 0.1 and a Top P value of 0.7.

### Manual validation

To evaluate human tracking performance against BehaveAgent, we designed a simple open field experiment. A single mouse was allowed to habituate to a 40 × 40 cm open field while its behavior was recorded using a GoPro Hero camera at 30 frames per second. For comparison, we used two tracking approaches: Coordinates generated by BehaveAgent with Molmo, and manual coordinate placement for each frame. Manual annotations were performed independently by two human subjects using a custom Python script with GUI. The interface allowed users to load videos and follow standardized instructions such as: “*Point precisely to the center of the white mouse*,” replicating the prompt given by BehaveAgent for consistent evaluation. To assess tracking accuracy, we computed pixel differences between human placed and Molmo generated coordinates across both the x and y axes. The resulting data were visualized using 3D trajectory plots (x, y position over time) to qualitatively compare the alignment of tracking paths between human subjects and BehaveAgent.

### Frontend

We implemented BehaveAgent’s frontend using Streamlit^36^. The sidebar provides direct video import functionality through a file uploader component supporting standard experimental formats (.mp4, .avi) with immediate preview rendering. The UI incorporates custom CSS theming through runtime style injection from themes folder. Analysis mode selection enables workflow customization based on user requirements. Media visualization uses a standardized pipeline supporting both video streams and static plots with automatic PDF conversion. User settings for filtering parameters (Kalman, median, Savitzky-Golay, VLM-based) are preserved through Streamlit’s session_state with nested configuration objects. We also implemented progress tracking with custom CSS classes for operational transparency during computation.

## Code and data availability

Code is available at http://github.com/LiuLab-Bioelectronics-Harvard/BehaveAgent

## Supporting information

Supplementary Video 2: BehaveAgent demo 1 - Novel object recognition.

Supplementary Video 3: BehaveAgent demo 2 - Drosophila courtship behavior.

Supplementary Video 4: BehaveAgent demo 3 - Primate object manipulation and recognition.

Supplementary Table 1: Video details.

## Acknowledgements

We acknowledge all the members of Liu Lab for their invaluable input during the creation of this manuscript and their support for this work. J.L. Acknowledge support from NIH/NICHD 1R01HD115272.

## Author contributions

A.A, Z.L. and J.L. conceived the idea. A.A and Z.L. developed the BehaveAgent method. W.W. and N.L. help implemented tools. X.Z. and R.L. prepared the behavior videos. B.C. and C.L. performed the human tracking analyses. A.A. and Z.L. prepared figures and drafted the initial manuscript. All authors contributed critical discussions and input on the figures and results. J.L. supervised the study.

## Extended Data Figures and Figure Captions

**Extended Data Fig. 1:**
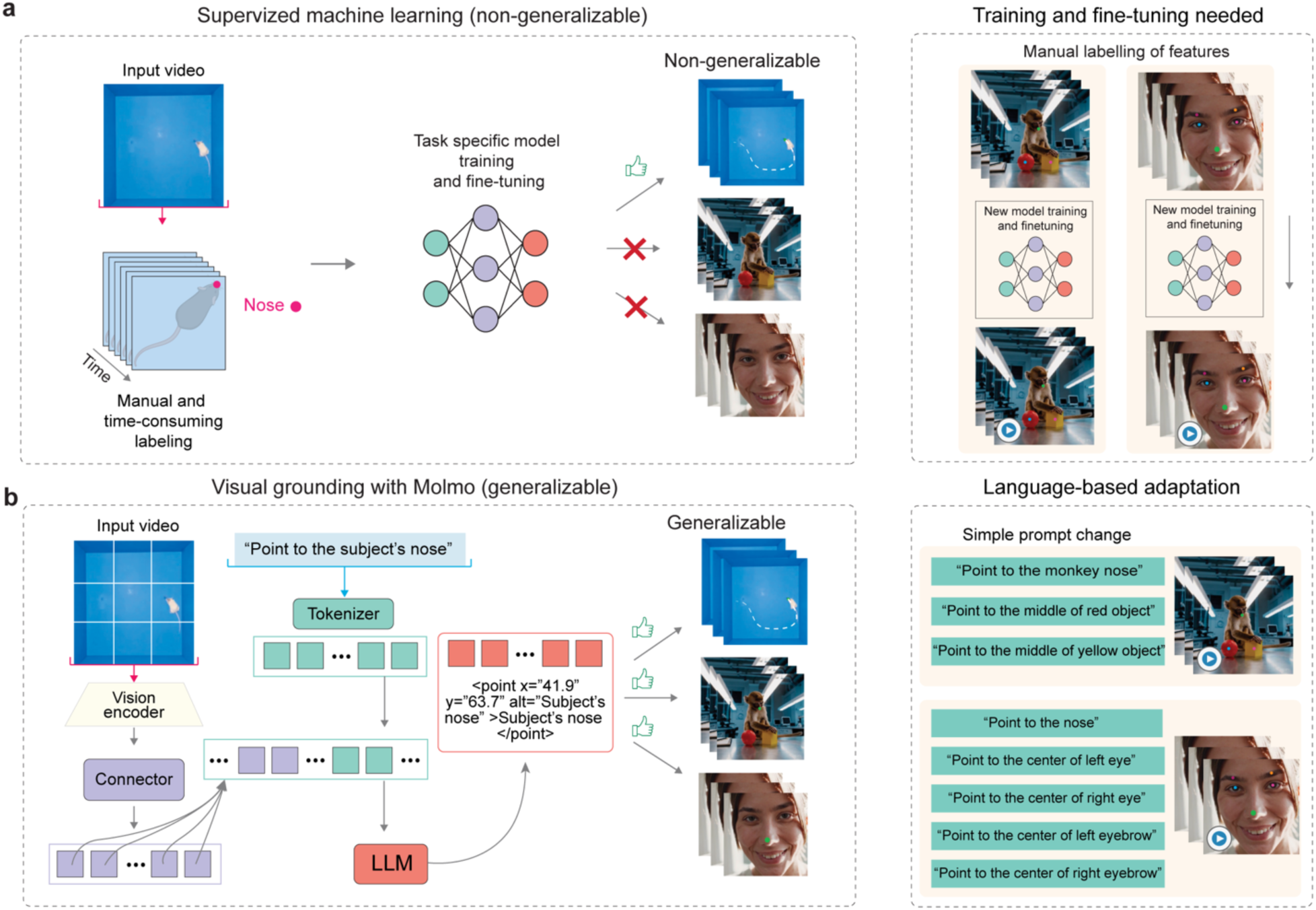
Comparison of traditional supervised machine learning and BehaveAgent for tracking and pose estimation. (**a**) Traditional supervised machine learning approaches for tracking and pose estimation requiring manual and time-consuming labeling of features, and task-specific model training and fine-tuning. These models lack generalizability across different species or experimental contexts, requiring new model training and fine-tuning for each new type of subject (rodent, primate, human) or behavioral paradigm. This process creates significant barriers to applying behavior analysis across diverse research contexts. (**b**) In contrast, BehaveAgent’s using visual grounding approaches such as Molmo that offers inherent generalizability without requiring any additional training. The Molmo system processes input videos through a vision encoder and connector architecture, which interfaces with a LLM to interpret natural language prompts. This architecture generates precise spatial coordinates that accurately locate the requested feature. The approach is easily adaptable to new inputs through simple prompt changes, allowing researchers to track different features across diverse subjects without requiring any new training data or model adjustments. This generalizability significantly enhances BehaveAgent’s ability to function across multiple species and behavioral contexts without specialized configuration.

**Extended Data Fig 2:**
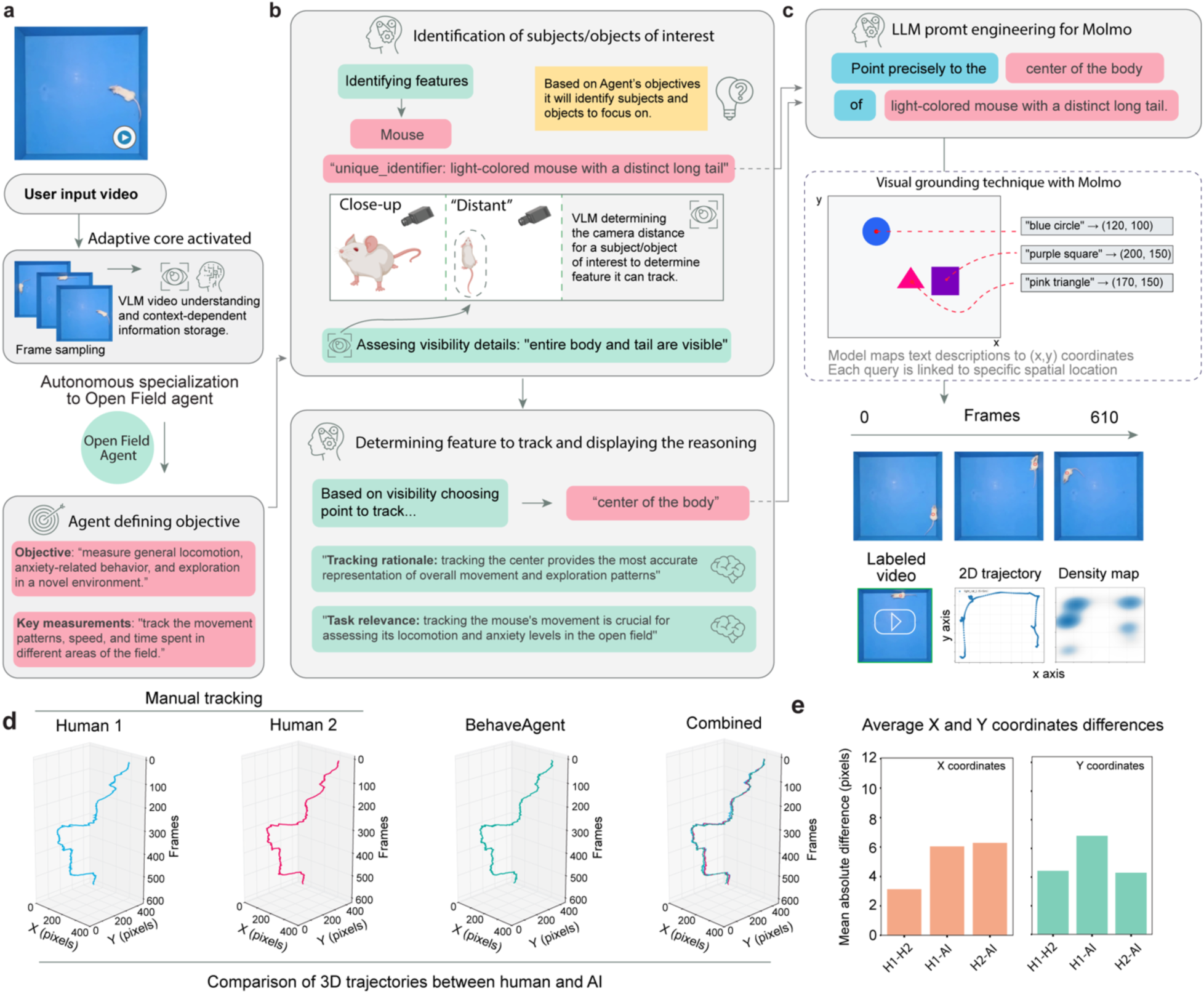
Comparison of 3D trajectories between human manual tracking and BehaveAgent. (a-c) BehaveAgent processing user input video by activating an adaptive core, automatically defining its objective, and transforming into an OpenField Agent tasked with tracking locomotion and anxiety-related behavior. Using VLM and LLM-guided reasoning, it identifies the subject, selects the optimal tracking point (“center of the body”), and employs visual grounding to extract consistent (x, y) coordinates and generate spatial tracking outputs. **(d)** Side-by-side comparison of 3D trajectory reconstructions from two human annotators and BehaveAgent, shown across x-y spatial dimensions over time (frames). The combined plot illustrates overall alignment between human (H1 and H2) and AI tracking. **(e)** Quantification of tracking accuracy using mean absolute differences in x and y coordinates across all frames.

**Extended Data Fig. 3:**
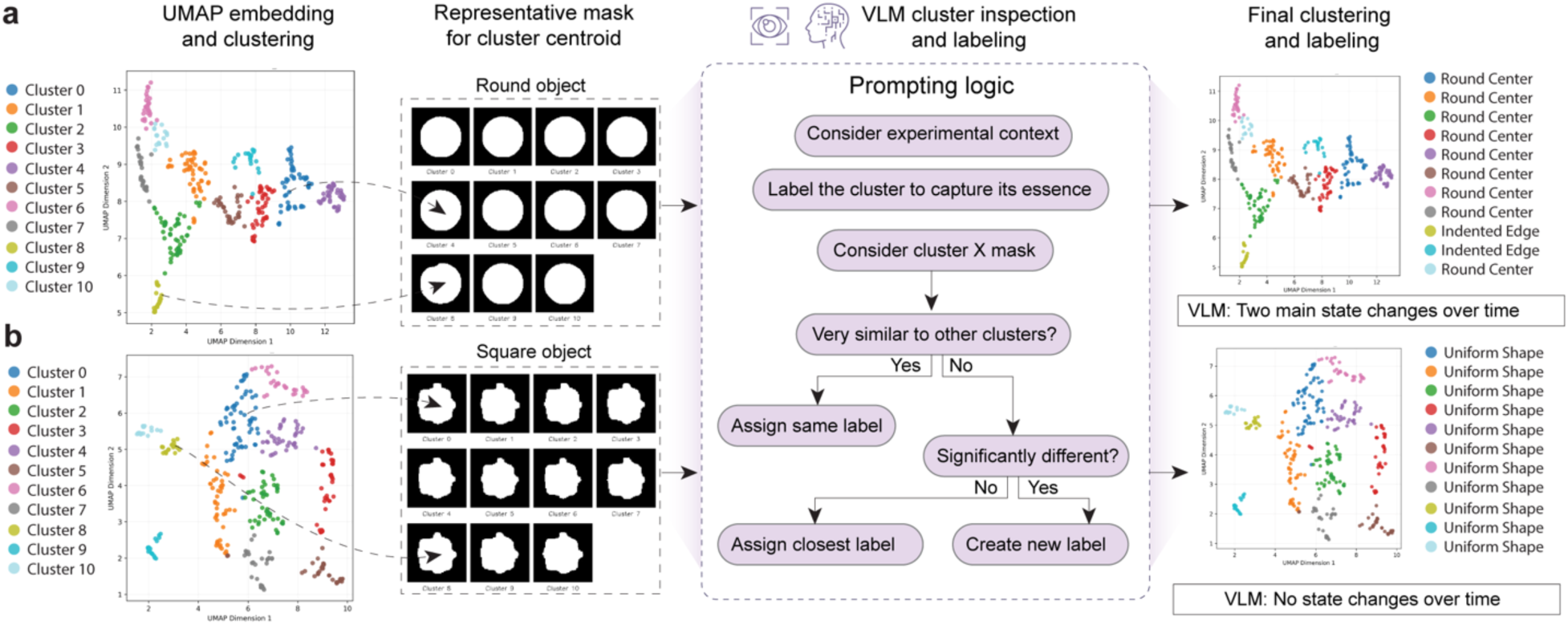
High-resolution object state clustering enables VLM-based refinement into behaviorally relevant labels. **(a)** UMAP embedding of round object mask features resulting in 11 visually separable clusters. Representative masks for each cluster centroid are shown, capturing fine differences such as roundness, indentation, or edge deformation. A VLM inspects each cluster mask and assigns semantic labels through a stepwise prompting logic that considers experimental context, cluster distinctiveness, and visual similarity. Final labeling reveals two dominant classes: “Round center” and “Indented edge,” indicating meaningful state transitions over time. **(b)** UMAP embedding of square object mask features producing 11 clusters, with representative masks showing minimal shape variation. VLM inspection confirms all clusters exhibit uniform structure. Final labeling assigns the same “Uniform shape” label to all clusters, and VLM interpretation concludes that no distinct state changes occurred over time for the square object.

**Extended Data Fig. 4:**
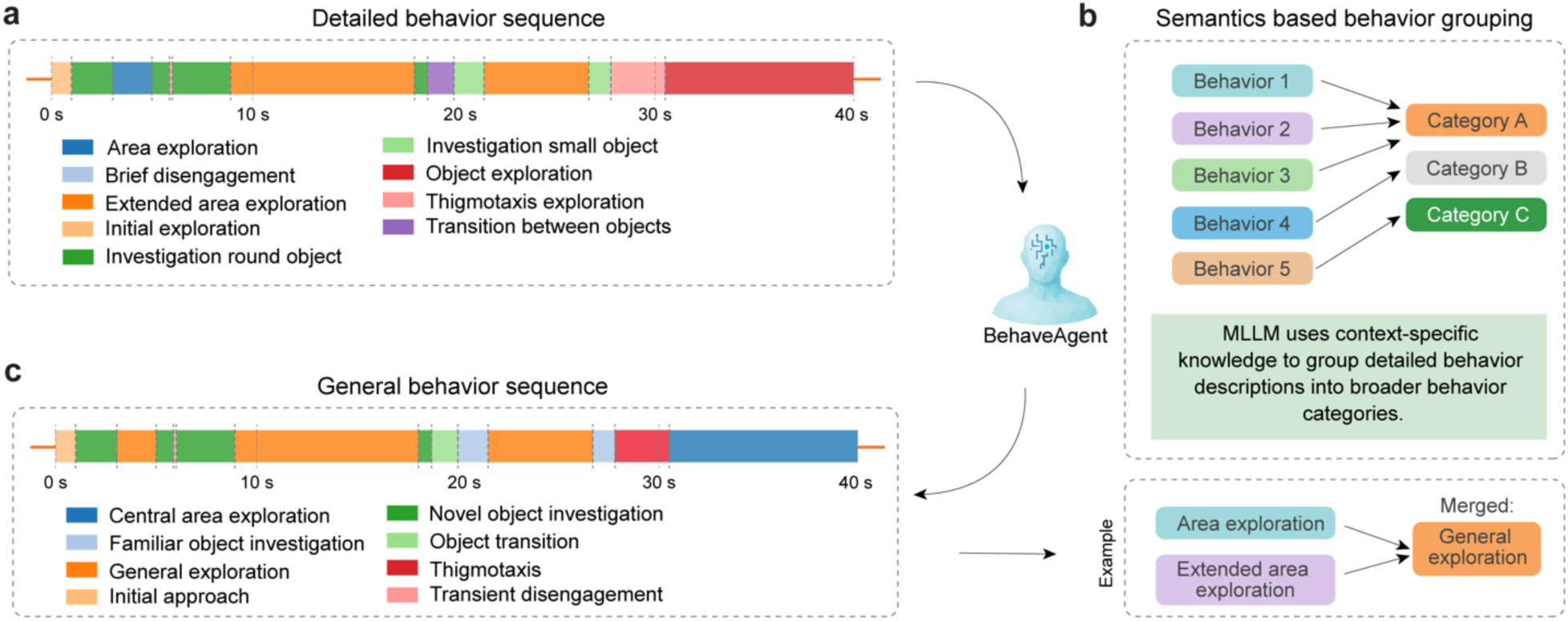
Merging detailed behavior descriptions into broader, semantically grouped categories. **(a)** BehaveAgent generating a fine-grained behavior sequence over the recording period. Detected states include diverse exploratory patterns and transitions, such as “Investigation small object,” “Thigmotaxis exploration,” and “Transition between objects.” **(b)** A multimodal LLM using context-specific semantic knowledge to group similar behavior segments into broader behavioral categories. Individual detailed behaviors (e.g., “Area exploration” and “Extended area exploration”) are merged into unified categories (e.g., “General exploration”) based on conceptual similarity. **(c)** The final general behavior sequence that reflects this abstraction, simplifying the behavioral timeline into a set of higher-level, interpretable categories such as “Novel object investigation,” “General exploration,” and “Thigmotaxis.”

**Extended Data Fig. 5:**
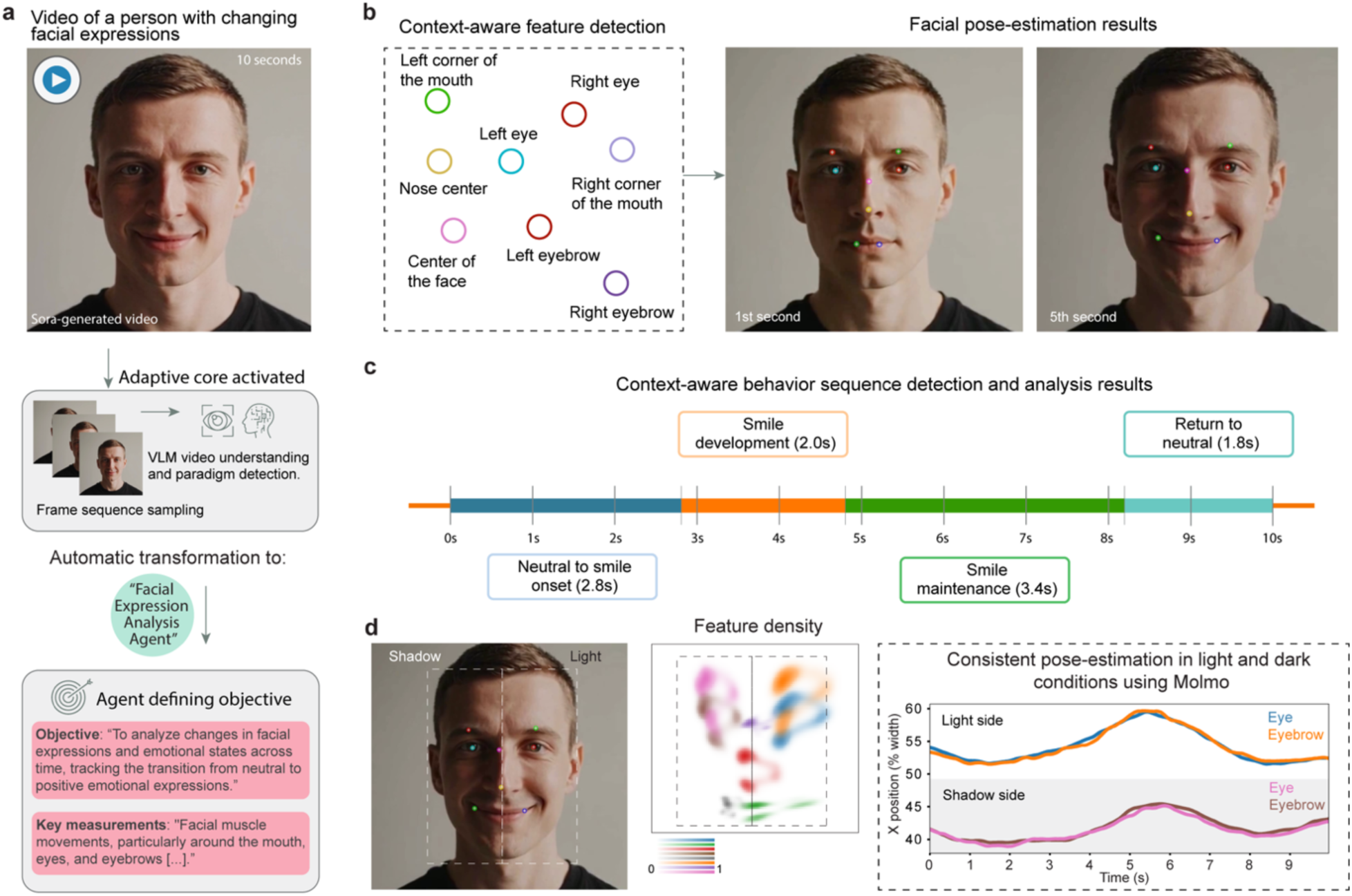
End-to-end human facial expression analysis. (**a**) An AI-generated video of a person displaying changing facial expressions is analyzed by BehaveAgent, which automatically transforms into a “Facial Expression Analysis Agent” with appropriate objectives and measurements. (**b**) The system performing context-aware feature detection of key facial landmarks (left panel), with results shown as pose-estimation of detected facial features (right panel). (**c**) BehaveAgent that segments the 10-second video into distinct emotional states on a timeline, identifying progression from neutral expression through smile development, maintenance, and return to neutral. (**d**) Results demonstrating robustness to challenging lighting conditions, maintaining consistent pose-estimation across shadowed and well-lit portions of the face. Sora-generated refers to the use of AI for video generation.

## Description of additional supplementary data

**Supplementary Video 1:** BehaveAgent autonomously detects experimental paradigms, defines task objectives, and performs tracking, segmentation, and behavior analysis across diverse species and contexts.

**Supplementary Video 2:** *BehaveAgent demo 1 - Novel object recognition*. (Sections of the videos with longer tool run times have been accelerated for demonstration purposes).

**Supplementary Video 3:** *BehaveAgent demo 2 - Drosophila courtship behavior*. (Sections of the videos with longer tool run times have been accelerated for demonstration purposes).

**Supplementary Video 4:** *BehaveAgent demo 3 - Primate object manipulation and recognition*. (Sections of the videos with longer tool run times have been accelerated for demonstration purposes).

**Supplementary Table 1:** Video details.

